# 16p11.2 Microduplication is Associated with Lobule-Specific Abnormalities in Cerebellar Structure and Function

**DOI:** 10.1101/2023.11.20.565320

**Authors:** Cessily Hayes, Hunter Halverson, Krisha Keeran, Krislen Tison, Kamilla Jacobo, Asriya Karki, Isaias Herring, Swetha Tunuguntla, Martha Pace, Binh Doan, Hsiang Wen, Annette Klomp, Marisol Lauffer, Marie E. Gaine, Krystal Parker, Aislinn J. Williams

**Author notes:** equally contributing authors (Krystal Parker, Aislinn J. Williams).

## Abstract

The 16p11.2 microduplication (16p11.2^dp/+^) is associated with several neuropsychiatric disorders including schizophrenia, autism spectrum disorder, bipolar disorder, intellectual disability, and attention deficit/hyperactivity disorder (ADHD). Cerebellar abnormalities have been increasingly implicated in these neuropsychiatric disorders, including those conferred by 16p11.2 microduplication. In 16p11.2^dp/+^ mouse models, the cerebellum is a site of transcriptional dysregulation, and cerebellar microcephaly has been reported in humans with 16p11.2 microduplication. Despite mounting evidence indicating a role for the cerebellum in neuropsychiatric disorders associated with this CNV, cerebellar cellular structure and cerebellar-dependent behavior in mice with 16p11.2 microduplication remain uncharacterized. To address this, we histologically labeled Purkinje cells (PCs) and molecular layer interneurons (MLIs) in a mouse model of 16p11.2 microduplication. We did not find any structural differences in cerebellar lobule IV/V, nor did we observe impairments in gait or motor coordination, behaviors that are associated with lobule IV/V. In contrast, we discovered a significant increase in calbindin/parvalbumin-positive PCs mislocalized to the granule layer of cerebellar lobule VI in 16p11.2^dp/+^ mice compared to wild-type (WT) littermates. Additionally, we found a significant decrease in parvalbumin-positive MLIs without a decrease in total DAPI-positive cell counts in lobule VI of 16p11.2^dp/+^ mice compared to WT littermates. Cerebellar lobule VI is associated with delay eyeblink conditioning, and 16p11.2^dp/+^ mice are impaired in cerebellum-dependent associative learning on this task. Specifically, 16p11.2^dp/+^ mice showed deficits in both conditioned response (CR) percentage and CR onset latency relative to WT mice. These results suggest that lobule VI–specific alterations to PC localization and MLI parvalbumin expression in 16p11.2^dp/+^ mice impair both cerebellar learning and adaptive timing of cerebellar-driven, conditioned responses. Thus, we have identified novel structural and functional alterations in the cerebellum that are associated with 16p11.2 microduplication. Importantly, individuals with schizophrenia and ADHD also show CR acquisition deficits in delay eyeblink conditioning. Together, these data suggest that the behavioral impairments in 16p11.2^dp/+^ mice resemble impairments seen in neuropsychiatric disorders linked to 16p11.2 microduplication in humans. Further investigation of cerebellar cortex neurons in 16p11.2^dp/+^ mice may provide insights into the pathogenesis of neuropsychiatric disorders linked to this copy number variant.

## INTRODUCTION

The World Health Organization estimates that 1 in 8 people worldwide live with a neuropsychiatric disorder (2019), yet the underlying mechanisms driving the ontogenesis of these conditions remains poorly understood. Certain copy number variants (CNVs) may increase the risk of developing one or multiple disorders and thus serve as a useful model to elucidate the shared etiology of neuropsychiatric disorders. Although rare in the general population, occurring in 1 of every 4216 live births [1], 16p11.2 microduplication has been associated with multiple neurodevelopmental and neuropsychiatric disorders, including schizophrenia (SCZ), autism spectrum disorder (ASD), bipolar disorder (BD), attention deficit hyperactivity disorder (ADHD), and intellectual disability (ID) [2–5, 6, 7–13]. Additionally, a genome-wide association study revealed that the 16p11.2^dp/+^ CNV conferred one of the highest genetic risks for the development of SCZ [14].

Although frontal-cortical abnormalities are well-established phenotypes of 16p11.2 microduplication [15–19] and neuropsychiatric disorders in general [20, 21], the cerebellum’s role in these disorders has been relatively underexplored. While the cerebellum has historically been considered a primary regulator of motor control, evidence from cerebellar stroke, lesion, and neuropsychiatric studies supports the cerebellum’s involvement in non-motor functions as well. Specifically, abnormalities in posterior cerebellar lobules (Lobule VI-X) contribute to alterations in executive function, linguistic processing, and spatial cognition, while motor alterations arise from abnormalities affecting the anterior lobules (Lobule I-IV/V) [22–31].

Furthermore, multiple behavioral studies in humans and mouse models have linked the cerebellum with neuropsychiatric disorders relevant to 16p11.2 microduplication. A commonly employed method to investigate cerebellar associative learning is delay eyeblink conditioning. Electrophysiological studies in animals have shown that the acquisition of conditioned responses relies on long-term depression at the parallel fiber-PC synapse and long-term potentiation at the mossy fiber-interpositus nucleus synapse [32–34], exemplifying the necessity of functional cerebellar circuitry for learning to occur on this task. Additionally, lesions in cerebellar lobule VI have been found to impair the learning and expression of conditioned responses in both humans and animals [35, 36]. Therefore, delay eyeblink conditioning serves as a useful measure of cerebellar integrity within the scope of neuropsychiatric disorders. Eyeblink conditioning studies in individuals with SCZ have revealed alterations to cerebellar-dependent associative learning, as they display reduced CR % [37–42] and longer CR onset latencies [40, 43], whereas children with ASD showed shorter CR onset latencies [44]. In animal models, the Erasmus Ladder provides a novel behavioral paradigm to evaluate cerebellar-dependent motor performance and learning within a single experiment. The cerebellar-dependent learning aspect of this task involves mice traversing a horizontal ladder, with the introduction of a perturbation paired with a tone cue. While initially unexpected, mice eventually learn to anticipate the perturbation and swiftly step to reach the goal box. It has been previously demonstrated that the inferior olive [45] and PCs are necessary for this learning process [46], underscoring the crucial role of the cerebellum in this paradigm. Moreover, experimental evidence links impairments in gait and motor coordination to lesions in cerebellar lobule IV/V in mice [26, 47]. *Shank* mouse models displaying phenotypes related to ASD exhibit cerebellar-mediated learning deficits without changes in motor performance on the Erasmus Ladder [48].

Along with alterations in cerebellar-dependent behavior, alterations in cerebellar structure have also been implicated in neuropsychiatric disorders, including those conferred by 16p11.2 microduplication. Histological examination in individuals with ASD has revealed reduced PC density and PC soma area in the cerebellar vermis and hemispheres [49–51], with some describing PC loss localized to cerebellar lobule VI and VII [52]. Reports are inconsistent in individuals with SCZ, with some describing PC abnormalities [53], and others finding no loss of PCs [54]. In the few studies that have examined the cerebellum in humans with 16p11.2 microduplication, smaller sizes of multiple brain structures, including the cerebellum, have been reported [55]. Duplicated genes in the 16p11.2 locus also appear to be more highly expressed in the cerebellum compared to other brain regions [56]. Given compelling evidence for structural and functional involvement of the cerebellum in neuropsychiatric disorders associated with 16p11.2 microduplication, there is a critical need for studies to comprehensively define its molecular, cellular, and functional significance.

Delineating the role of the cerebellum in this CNV can be achieved through the assessment of animal models, such as the Horev et al. 16p11.2 microduplication mouse model [57] utilized in this study. Various structural and behavioral phenotypes have been reported in this mouse model previously, although the cerebellum was not a major point of focus for most studies. Structurally, 16p11.2^dp/+^ mice display reduced size of basal forebrain, hypothalamus, medial septum, and periaqueductal gray [57], mirroring the microcephaly phenotype in individuals with 16p11.2 microduplication [55]. Interestingly, a bulk RNA sequencing study in 16p11.2^dp/+^ mice revealed the highest occurrence of differentially expressed genes in the cerebellum compared to other brain regions [58]. Behaviorally, 16p11.2^dp/+^ mice do not display motor coordination deficits [59], but were found to exhibit hypolocomotion, social and cognitive deficits, and increased repetitive self-grooming [19, 57, 59]. However, there are conflicting findings for prepulse inhibition of the startle response [19, 59]. Prepulse inhibition (PPI) is a phenomenon in which a weak stimulus (prepulse) can suppress a natural startle response to a subsequent stronger startle stimulus (pulse). This task is used to assess sensorimotor gating, the process of filtering irrelevant sensory information, as deficits in attenuation of the startle response are common in individuals with SCZ, ASD, and ADHD [60–62]. While some studies report no PPI impairments in 16p11.2^dp/+^ mice [59], others have reported PPI deficits specific to female 16p11.2^dp/+^ mice [19]. Considering the high association of SCZ with this CNV, and the robust impairments in PPI reported in individuals with SCZ, it is surprising that there are conflicting findings regarding impairments in PPI in 16p11.2^dp/+^ mice.

Given inconsistent findings for PPI and a clear, yet undefined, role for the cerebellum in the pathophysiology of neuropsychiatric disorders that involve 16p11.2 microduplication, there is a great opportunity to elucidate the structure and function of the cerebellum in mice with this CNV. Here, we analyze cerebellar cellular composition, cerebellar-dependent motor function, associative learning, and sensorimotor gating. Although our data do not support gross cerebellar structural alterations in 16p11.2^dp/+^ mice, we discovered lobule VI-specific differences with respect to PC localization and reduced parvalbumin labeling in apical MLIs without a concomitant reduction in total molecular layer cells in 16p11.2^dp/+^ mice. Behaviorally, there were deficits in cerebellar associative learning during delay eyeblink conditioning and deficits in sensorimotor gaiting in female 16p11.2^dp.+^ mice assessed by PPI, with no alterations to gait or motor coordination assessed by the Erasmus Ladder. Together, our work further supports cerebellar involvement in the pathogenesis of neuropsychiatric disorders associated with 16p11.2 microduplication.

## MATERIALS AND METHODS

### Animals

Mice carrying a heterozygous duplication of the 7F4 chromosomal region corresponding to human 16p11.2 microduplication (16p11.2^dp/+^ mice) were recovered from cryopreserved embryos by The Jackson Laboratory (Strain #:013129) [57]. Breeding pairs of 16p11.2^dp/+^ mice were maintained on a C57BL/6J x129S1/SvImJ (referred to as B6129SF1/J) hybrid background (Jax Strain #:101043) by crossing 16p11.2^dp/+^ offspring with B6129SF1/J WT mice purchased from Jackson Laboratory (Bar Harbor, Maine). To generate experimental animals, 16p11.2^dp/+^ mice were bred to WT mice to produce litters with combinations of 16p11.2^dp/+^ and WT offspring. All mice were adults (at least 10 weeks old) at the time of testing. Sample sizes and ages of mice are indicated in each figure. Littermates were same-sex group housed under regular light cycle on/off at 0900/2100 DST (0800/2000 non-DST). The average ambient temperature was 22°C and mice were provided with food and water ad libitum, with enriched paper bedding used in all homecages. Each experiment was conducted during the animals’ light cycle and equipment was cleaned between every trial/session with 70% ethanol.

All experiments were carried out in a manner to minimize pain and discomfort, and animals were monitored after each experiment to ensure their health and safety. All experiments were conducted according to the National Institute of Health guidelines for animal care and were approved by the Institutional Animal Care and Use Committee at the University of Iowa.

### Histological examination

Mice were anesthetized by intraperitoneal injections of 17.5 mg/ml Ketamine/ 2.5 mg/ml Xylazine at a dose of 0.1 ml per 20 g. Cardiac perfusions were performed with ice-cold 0.1 M phosphate-buffer (PB; pH 7.4) followed by 4% paraformaldehyde (PFA) diluted in PB. Whole brains were dissected and stored in 4% PFA for at least 24 hours, then cryoprotected in 30% sucrose diluted in PB for approximately 72 hours. Brains were rinsed with PB and frozen in optimal cutting temperature compound using dry ice in 2-methylbutane. Brains were stored at –80°C for at least 24 hours before being serially sectioned sagittally through the cerebellar vermis at 20 μm, on a cryostat set at –20°C. We retained every 5^th^ section; therefore, serial sections were approximately 100 μm apart. Sections were air dried and stored at –20°C until they were stained.

### Immunohistochemistry

Four sections from each mouse were stained using primary antibodies calbindin D28K anti-rabbit recombinant polyclonal (1:200; ThermoFisher) and parvalbumin anti-mouse monoclonal (1:200; Swant). The primary antibodies were diluted in blocking buffer made from 5% normal donkey serum and 0.1% Triton-X and incubated on slides in a humidified chamber at 4°C overnight. Sections were then labeled with secondary antibodies Donkey anti-rabbit 488 (1:500; Jackson), Donkey anti-mouse 594 (1:500; Jackson), and DAPI solution (1:1000; ThermoFisher). Diluted secondary antibodies and DAPI solution were incubated at room temperature for two hours in a humidified chamber. Sections were then coverslipped with Prolong Diamond Antifade Mountant (Invitrogen).

### Nissl Staining

Two sections per mouse were stained with thionin. Sagittal cerebellar vermis sections were first rehydrated by dipping slides into deionized (DI) H_2_O for 5 seconds. Excess water was drained off slides before dipping slides up and down gently in thionin stain for approximately 90 seconds. Slides were then rinsed briefly in DI H_2_O before placing into a 70% alcohol solution for 5 minutes and then a 95% alcohol solution for 5 minutes. Slides were then placed in a 95% alcohol/acetic acid solution until desired color was achieved. Afterwards, slides were placed in a 100% alcohol solution for 5 minutes before removing and placing into a separate 100% alcohol solution for another 5 minutes. Slides were then moved through three separate citrus clearing solvent baths for 5 minutes each. After, slides were coverslipped with Permount.

### Ectopic Purkinje Cell Counts

Four immuno-labeled sagittal sections per mouse as described above were imaged at 20X in three locations (anterior, medial, and posterior) of lobule IV/V and lobule VI of the cerebellar vermis on an Olympus IX83 fluorescence microscope for the analysis of ectopic PCs. Twelve images per mouse (3 images/lobule/4 sections/mouse) were used. Counting was performed by raters blinded to genotype. Calbindin+ cells in the granule layer of cerebellar lobule IV/V and VI were classified as an ectopic PC if they met the following criteria: contained a nucleus, cell diameter between 45 – 80 µm, calbindin/parvalbumin immunoreactivity in the entire cell or the membrane, and the calbindin immunoreactivity of the ectopic PC must be distinct from background fluorescence. Ectopic PCs were counted manually in ImageJ for each image and recorded for both lobule IV/V and VI. Ectopic PC counts were added together within a section and resulted in a total of four data points per mouse.

### Typically Located Purkinje Cell Cross-Sectional Soma Area

One thionin-stained sagittal section per mouse was imaged at 10X in lobule VI of the cerebellar vermis using bright field on an Olympus IX83 fluorescence microscope for the analysis of typically located PC soma cross-sectional area. We used thionin staining to quantify typically located PC soma cross-sectional area because calbindin labeling did not reliably fill the somata of typically located PCs, particularly in the portions of lobule VI adjacent to lobule IV/V. Typically located PC soma cross-sectional area measurements were performed in ImageJ. First, in ImageJ, the image scale was calibrated by measuring the length of the scale bar (known distance) to convert distance measured in pixels to microns. In total, we measured 24 PC somata per mouse. We selected six consistent locations within lobule VI to measure typically located PC soma cross-sectional area, selecting 4 PCs to measure from in each area to consider variability in soma size across the lobule. A PC’s soma cross-sectional area was only quantified if the PC displayed a primary proximal dendrite in attempt to capture the maximal cross-sectional area of a PC. A PC’s soma cross-sectional area was measured by drawing a polygon around the entire soma in ImageJ. The polygon area was then measured and recorded in units of µm^2^. Measurements were performed by raters blinded to genotype. The 24 typically located PC somata cross-sectional area measurements were averaged within a mouse and resulted in one data point per mouse. Additionally, typically located PC somata cross-sectional areas were measured in ImageJ using confocal microscopy to verify that typically located PC somata cross-sectional areas measured via thionin were not significantly different from the same typically located PC somata cross-sectional areas measured via calbindin on the confocal.

### Ectopic Purkinje Cell Cross-Sectional Soma Area

Using the same immuno-labeled sagittal sections from ectopic PC counts, three ectopic PCs in the granule layer of lobule VI per 16p11.2^dp/+^ mouse were imaged at 40X with a zoom factor of 1.5 and resolution of 2048 x 2048 on a Leica SPE Confocal microscope. Thionin staining was not used to quantify ectopic PC soma cross-sectional area because ectopic PCs in the granule layer were indistinguishable from granule cells by thionin staining. Additionally, a confocal microscope was used to capture ectopic PCs instead of an epifluorescent microscope to reduce light scattering, which artificially inflates calbindin-labeled PC soma cross-sectional area. Ectopic PC soma cross-sectional area measurements were performed in ImageJ. First, the image scale was calibrated and the cross-sectional area of three ectopic PC somata previously identified in 16p11.2^dp/+^ mice were measured by drawing a polygon around the entire soma in ImageJ. The polygon area was then measured and recorded in units of µm^2^. The three ectopic PC soma cross-sectional area measurements were averaged within a mouse and resulted in one data point per mouse.

### Typically Located Purkinje Cell Density

The same immuno-labeled sagittal sections from ectopic PC counts were imaged at 4X to capture the entirety of lobule IV/V and lobule VI on an Olympus IX83 fluorescence microscope for the analysis of typically located PC density. Cell counts were performed in ImageJ by raters blinded to genotype. First, the image scale was calibrated. Typically located PC counts were performed by adding a line in ImageJ across the PC monolayer, acquiring the distance of line in microns, and counting the number of PCs that spanned the line. This method was repeated in five consistent locations in lobule IV/V and lobule VI for each mouse section. In some sections, the PC somata were not reliably filled by calbindin in portions of lobule VI adjacent to lobule IV/V. To ensure accurate cell counts in these areas, we used thionin stained sections. The cell counts were divided by the length of the line in microns to acquire a density measurement. Each of the five typically located PC density measurements were averaged to obtain one value per mouse.

### Molecular Layer Interneuron Counts

One immuno-labeled sagittal section per mouse was imaged at 4X to capture the entirety of lobule IV/V and lobule VI in the cerebellar vermis on an Olympus IX83 fluorescence microscope for MLI cell counts. Cell counts were performed in ImageJ. First, in ImageJ, the image scale was calibrated and the DAPI and parvalbumin channel images (1 image and 1 section per mouse) were manually thresholded in ImageJ. The purpose of thresholding was to create a binary, such that DAPI+ and parvalbumin+ cells could be quantified in a semi-automated fashion. By assigning background fluorescence as a value of “0” and DAPI+ or parvalbumin+ cells assigned a fluorescence value of “1”, ImageJ quantified pixels with a value of “1” as a DAPI+ or parvalbumin+ cell. After thresholding, polygons were drawn in the molecular layer using the DAPI channel and copied into the parvalbumin channel to maintain the same polygon size. The polygon area was then measured and recorded in units of µm^2^. The ImageJ “analyze particles” function was used to count both DAPI+ and Parvalbumin+ cells within the polygon, using the following parameters: size: 2-infinity (pixel units) and circularity: 0.00-1.00. This created regions of interest for each cell, and cell counts were recorded for both channels. DAPI+ and Parvalbumin+ cell counts were converted into units of cells per 0.1 mm^2^ using the following equation:

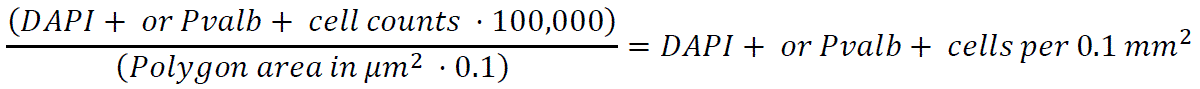

Thresholding and analysis were performed by raters blinded to genotype. MLI counts were performed for both lobule IV/V and VI. Five polygons per lobule in consistent locations across each mouse section were used to capture the entire molecular layer of each lobule. The cell counts for each polygon were then averaged and resulted in one value per mouse.

To determine if the distribution of DAPI+/parvalbumin+ MLIs was altered based on their locations within the molecular layer, we used a modified version of the procedure described above. Instead of drawing polygons spanning the entire width of the molecular layer, the thickness of the molecular layer of lobule VI was measured in ImageJ and the length divided to create inner 1/3 and outer 2/3 polygons (ten polygons in total per mouse). We used this anatomical distinction to operationally define and distinguish between early and late-born MLIs. Early born MLIs reside largely in the inner 1/3 of the molecular layer (defined as basal 1/3 MLIs), while later-born MLIs reside mostly in the outer 2/3 of the molecular layer (defined as apical 2/3 MLIs) [65].

### Behavioral procedures

#### Erasmus Ladder

The Erasmus Ladder task (Noldus, Wageningen, The Netherlands) is a test of motor function and motor associative learning that relies on the cerebellum [45]. It is made up of two enclosed boxes separated by a ladder, with a row of high rungs for the animals to efficiently cross the ladder, and a second row of lower rungs that record missteps in between the high rungs. In the first four days of training, animals are tested for abnormalities in gait adaptation, motor coordination, and associative learning. In the last four days of training, mice are tested for abnormalities in tone-cued cerebellar dependent associative motor learning. Prior to Erasmus Ladder, all mice used in the behavior cohorts were weighed.

Mice were trained to traverse the Erasmus Ladder for 42 trials per day for four consecutive days. Trials consisted of one run across the ladder into the goal box and were separated by a random inter-trial interval ranging from 11-20 seconds. In the first four days of training, a bright light cue indicated the start of a trial. Animals that did not leave the starting box following 3 seconds of light cue stimulus experienced a tailwind of pressurized air to motivate them to cross the ladder. The apparatus measured the animal’s steps across a discontinuous ladder and detected errors for each trial. Data were analyzed for the following: number of trials the animal left on light cue, number of trials the animal left on air cue, number of trials the animal went onto the ladder before a cue was given, frequency of the animal to return to the goal box, number of trials the animals refused to leave goalbox, percentage of missteps (lower rung steps/steps on ladder), percentage of correct long steps (skipping at least one high rung between steps), percentage of correct short steps (using consecutive high rungs). Post-perturbation step time was used to assess the baseline step time of an animal.

On the last four days of training, mice are tested for abnormalities in tone-cued cerebellar dependent associative motor learning. Mice were conditioned using a 75 decibel (dB) tone (CS) to anticipate the elevation of a ladder rung (US), also called the perturbation, in which the mice had to learn to time their jump in accordance with the auditory cue. Data were analyzed for time the animal spent between steps immediately preceding the perturbation (pre-perturbation) and steps immediately following the perturbation (post-perturbation) for the last 5-8 days of the task. Learning is shown when animals spend less time on the pre and post perturbation step following consecutive days of training.

#### Eyeblink Conditioning

Delay eyeblink conditioning is a classical conditioning paradigm where eyelid closure, as an unconditioned response, initially occurs after the onset of the unconditioned stimulus (an air puff to the eye). Through repeated exposures to a conditioned stimulus (a light) and the air puff, adaptively timed eyelid closure begins to coincide with the light conditioned stimulus, resulting in a conditioned eyeblink response.

Prior to eyeblink conditioning, adult mice were anesthetized with isoflurane and placed into a stereotaxic frame. A heating pad was used to maintain body temperature during surgery. After the onset of anesthesia, 1 mg/mL of meloxicam per 20 g of the animal’s weight was administered for post-operative analgesia. The head was leveled and a midline incision was made to expose the skull. The skin and underlying fascia was removed with tweezers and a cotton tipped applicator soaked in 70% ethanol. Once the skin was removed and the skull was dry, two small holes were drilled ∼2 mm caudal and lateral to bregma. Self-tapping machine screws were secured to the skull before centering a stainless steel headplate between the screws. The screws and headplate were fixed to the skull with Metabond cement (Parkell, Inc). After the cement dried, mice were returned to their home cage to recover under a heat lamp.

Before the start of an eyeblink conditioning session, each mouse was lightly anesthetized with isoflurane in an induction chamber. All eyeblink sessions were conducted inside an operant chamber with white noise (65 dB) generated through a speaker. After the onset of anesthesia, the headplate was fastened to a pair of machined rods with 2-56 machine screws. Each rod was attached to posts that surrounded a low friction freely moving Styrofoam cylinder. The height of the machined rods was adjusted so that each mouse could walk freely on the cylinder after the headplate was secured. A high-speed (200 frames/s) monochrome camera (Allied Vision or Basler) and infrared illumination were attached to a different post and were directed toward the right eye. For the air puff unconditioned stimulus (US) (20-30 PSI, 20-30 ms), plastic tubing connected to an air compressor/regulator was attached to a 23-gauge needle housed in an adjustable case. The 23-gauge needle was adjusted so the final position was 3 mm from the right eye and directed at the middle of the cornea. The intensity of the air puff unconditioned stimulus (US) was 6-8 PSI measured at the end of the needle. Time of actual puff delivery was corrected to account for the delays introduced by the solenoid and tubing, as measured at the end of the needle by a silicon pressure sensor (Honeywell HSCSANN 150PG2A3, DigiKey). The conditioned stimulus (CS) was a blue light LED mounted on a post in front of the mouse and the timing of the CS and US was controlled by a custom Arduino-based device [66].

Mice received two habituation sessions prior to eyeblink conditioning. Before the habituation (and eyeblink conditioning) sessions, mice were initially lightly anesthetized with isoflurane in an induction chamber. After breathing slowed, they were transferred to the operant chamber and secured over the cylinder. The habituation sessions allowed mice to move freely on the cylinder while head-fixed for one hour without presentation of the CS or US. Following habituation, mice were given ten daily eyeblink conditioning sessions (up to 100 trials per day) consisting of 10 trial blocks (9 CS/US paired, 1 CS alone trial) with an interstimulus interval of 400 ms. After mice were secured over the cylinder and had recovered from anesthesia, they received presentations of the air puff US to calibrate the reflexive eyelid closure. Eyelid movements were measured frame-by-frame by calculating the area of the eye visible within a custom fit elliptical region of interest. Raw pixel counts were normalized into units of fraction of eyelid closure (FEC) ranging from fully open (0) to fully closed (1). Once full eyelid closure was calibrated the eyeblink conditioning session was started.

Data were organized and analyzed in MATLAB using custom written software. Eyelid traces for each trial were extracted from the high-speed video files using the “pixel area” (FEC) algorithm. For paired trials, a conditioned response (CR) was operationally defined as an increase in eyelid amplitude during the final 300 ms of the CS that was greater than 10% of the unconditioned response amplitude to the air puff (full eyelid closure) for each trial. For CS alone trials, a CR was operationally defined as an increase in eyelid amplitude 100 ms after CS onset that was greater than 10% of the session average unconditioned response amplitude. CR onset latency was defined as the first time point when eyelid amplitude reached 10% of the unconditioned response amplitude for a given trial (paired trials) or 10% of the session average unconditioned response (CS alone trials). CR maximum amplitude was defined as the largest eyelid amplitude value during the interstimulus interval. Data from the final 300 ms of the CS was used for CR onset latency and amplitude analysis to limit data to learned responses that could be driven by cerebellar output.

#### Prepulse Inhibition

Mice were placed in an isolation cabinet and restrained in a clear acrylic cylinder. The tremble response of the animal was measured via an accelerometer underneath the chamber and the testing apparatus consisted of a startle response box (SR-LAB, San Diego Instruments). Mice were habituated to the testing chamber for 10 minutes with a consistent background white noise level of 65 dB, which continued for the duration of the experiment. Each mouse underwent 64 trials. The first and last 6 trials (Block I and Block III) consisted of solely pulse alone trials to verify that mice were not acclimating to the startle pulse over the experiment. All trials were presented with a randomly spaced inter-trial interval ranging from 7 to 15 seconds. The startle pulse was set to 120 dB and the prepulse intensities were set to 5, 10, and 15 dB above the background noise in the testing room. The startle response was recorded in millivolts in SR-LAB software. Percent PPI was calculated by normalizing the startle response of the pulse alone trials from block II using the following equation:

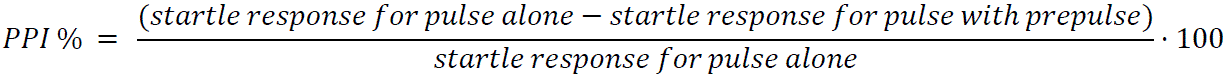

#### Statistics

Histological data were analyzed and graphed in GraphPad Prism 9. The Shapiro-Wilk normality test was applied to all data sets. A nested t test was used to detect group differences for ectopic PC counts. A nonparametric, unpaired t test (Mann-Whitney test) was used to detect group differences for typically located PC soma cross-sectional area. An unpaired t test was used to detect group differences for ectopic PC soma cross-sectional area. A two-way ANOVA was used to detect group differences for typically located PC density. Multiple unpaired t tests using the Holm-Sidak method were used to compare DAPI+ and parvalbumin+ MLI cell counts. In lobule VI, multiple unpaired t tests using the Holm-Sidak method were used to compare DAPI+ basal 1/3, DAPI+ apical 1/3, parvalbumin+ basal 1/3, and parvalbumin+ apical 2/3 MLI cell counts in WT and 16p11.2^dp/+^ mice.

Erasmus Ladder data were analyzed via linear mixed-effects modeling in RStudio with the ImerTest and emmeans packages and graphed in GraphPad Prism 9.1. Eyeblink conditioning data were analyzed using custom-written codes in MATLAB (MathWorks) and repeated-measures ANOVAs were done using SPSS. Group data was graphed in MATLAB and edited using Adobe Illustrator. Fisher’s Least Significant Difference (LSD) was used for post-hoc tests. All PPI data were analyzed to assess for sex, genotype, decibel, and block differences. Normality tests were applied to all data sets. Data were graphed and analyzed using GraphPad Prism and R (R4.1.1, emmeans 1.7.0, lme4 1.1.27.1, lmerTest 3.1-3). Data are graphically represented as mean ± standard error of the mean (SEM) for each group. Results were considered significant when *p* < 0.05 (denoted in all graphs as follows: \**p* < 0.05; \*\**p* < 0.01).

## RESULTS

### Histological Examination

#### Purkinje Cells

##### 16p11.2^dp/+^ mice display a significant increase in calbindin/parvalbumin+ ectopic Purkinje cells in the granule layer of lobule VI

Given evidence of cerebellar microcephaly in humans with 16p11.2 microduplication [55], we first sought to determine if there were structural alterations in the cerebellum of 16p11.2^dp/+^ mice. We observed expected overall cerebellar foliation and structure, including a granule cell layer, PC layer, and molecular layer. However, we discovered calbindin/parvalbumin+ PCs throughout the granule layer of multiple cerebellar lobules in both 16p11.2^dp/+^ mice and WT littermates. Therefore, we quantified ectopic PCs in both cerebellar lobule IV/V (anterior cerebellum) and lobule VI (posterior cerebellum). No differences were found in the number of calbindin/parvalbumin+ ectopic PCs localized to the granule layer of lobule IV/V between WT and 16p11.2^dp/+^ mice (nested t test, p = 0.3974; WT mean = 4.850, 16p11.2^dp/+^ mean = 6.800) (**Fig. 1A**). Conversely, we found a significant increase in the number of calbindin/parvalbumin+ ectopic PCs in the granule layer of cerebellar lobule VI in 16p11.2^dp/+^ mice relative to WT littermates (nested t test, p = 0.0036; WT mean = 2.550, 16p11.2^dp/+^ mean = 9.000) (**Fig. 1B**). We anecdotally observed that ectopic PCs were more frequent in posterior compared to anterior lobules (data not shown), and they appear to have consistently inelaborate dendritic arbors (**Fig. 1C and 1D**). Ectopic PCs appeared less intensely labeled for calbindin and parvalbumin than typically localized PCs, although they did express both PC markers (**Fig. 1E and 1F**).

**Figure 1.**
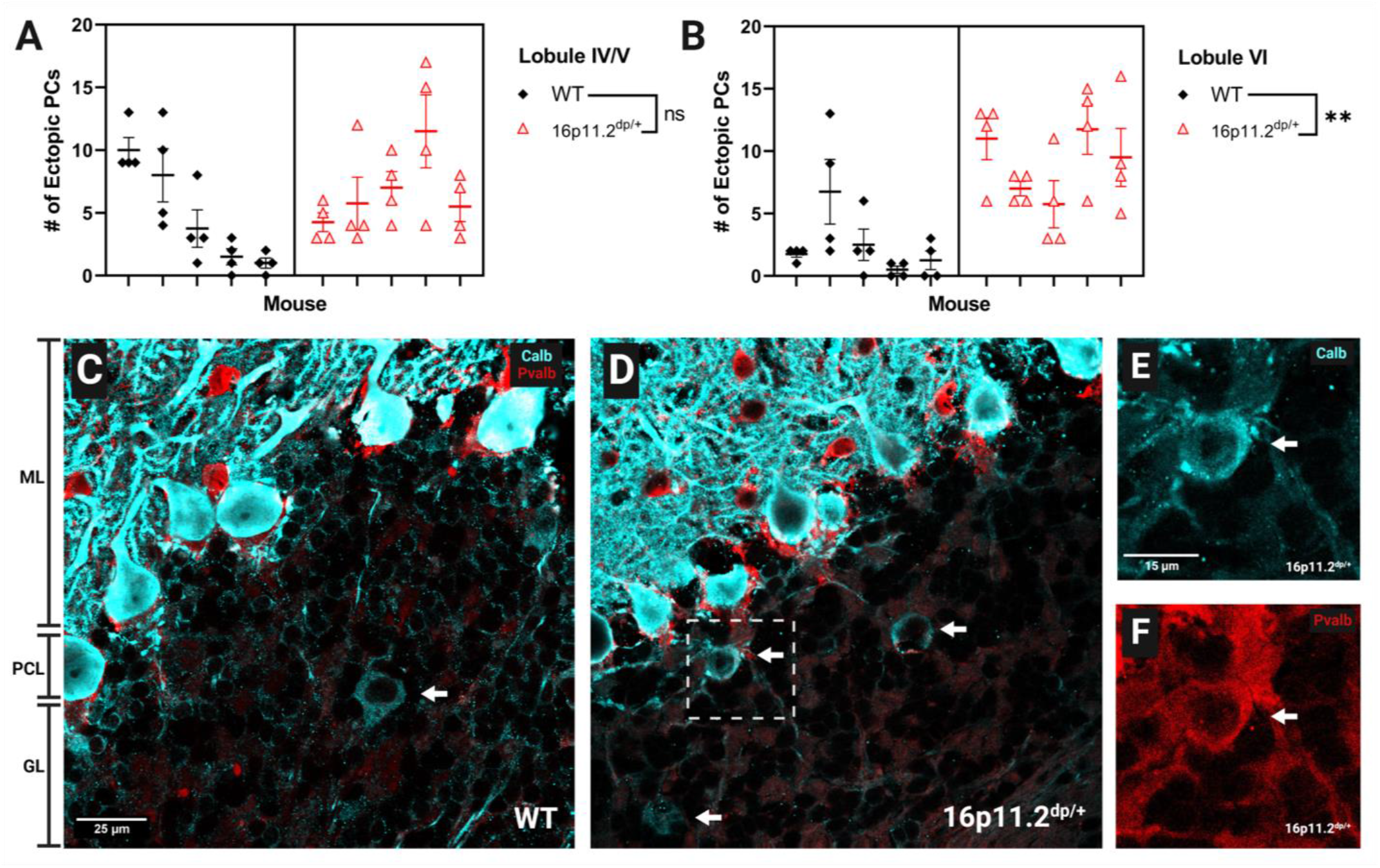
Lobule VI-specific increase in calbindin/parvalbumin+ ectopic Purkinje cells in 16p11.2^dp/+^ mice. **A.** No differences were found in the number of ectopic Purkinje cells (PCs) in lobule IV/V. **B.** A significant increase in ectopic PCs located in the granule layer of lobule VI was found in 16p11.2^dp/+^ mice compared to WT littermates. **A-B:** Mean with SEM, n = 10 (WTs: n = 3 females, n = 2 males; 16p11.2^dp/+^: n = 2 females, n = 3 males), mice were between 3.3-3.5 months old at the time of the experiment. **C.** Representative 40X confocal image (zoom factor 1.5) of calbindin (calb), shown in cyan, and parvalbumin (pvalb), shown in red, immunostaining of lobule VI in WT tissue. White arrow indicates calbindin/parvalbumin+ ectopic PC located in the granule layer. Cerebellar layers are indicated as molecular layer (ML), Purkinje cell layer (PCL), and granule layer (GL). **D.** Representative 40X confocal image (zoom factor 1.5) of calbindin/parvalbumin immunostaining of lobule VI in 16p11.2^dp/+^ mice. White arrows indicate calbindin/parvalbumin+ ectopic PCs located in the granule layer. **E and F.** Close up of ectopic PC in white box from **D** exhibiting both calbindin (**E)** and parvalbumin immunoreactivity **(F).**

##### No differences in the density or soma size of typically located Purkinje cells in 16p11.2^dp/+^ mice

Given the significant increase of ectopic PCs in the granule layer of lobule VI in 16p11.2^dp/+^ mice, and the previous findings of decreased PC density and PC soma area in humans with neuropsychiatric disorders associated with 16p11.2 microduplication [49–53], we sought to examine if alterations to cell counts or cell morphological features were present in typically located PCs. To determine if ectopic PCs resulted in depleted density of typically located PCs in lobule IV/V or VI, we investigated typically located PC density. We found no differences in typically located PC density between WT and 16p11.2^dp/+^ mice (2-way ANOVA, no main effect of genotype, p = 0.2374). However, there was a main effect of lobule, such that the PC density in lobule IV/V was less than lobule VI (2-way ANOVA, main effect of lobule, p = 0.0306) (**Fig. 2A**). Morphologically, no differences were found in lobule VI 16p11.2^dp/+^ typically located PC soma cross-sectional area relative to WT littermates (Mann-Whitney test, p = 0.7937; WT mean = 245.0 µm^2^, 16p11.2^dp/+^ mean = 236.8 µm^2^) (**Fig. 2B**).

**Figure 2.**
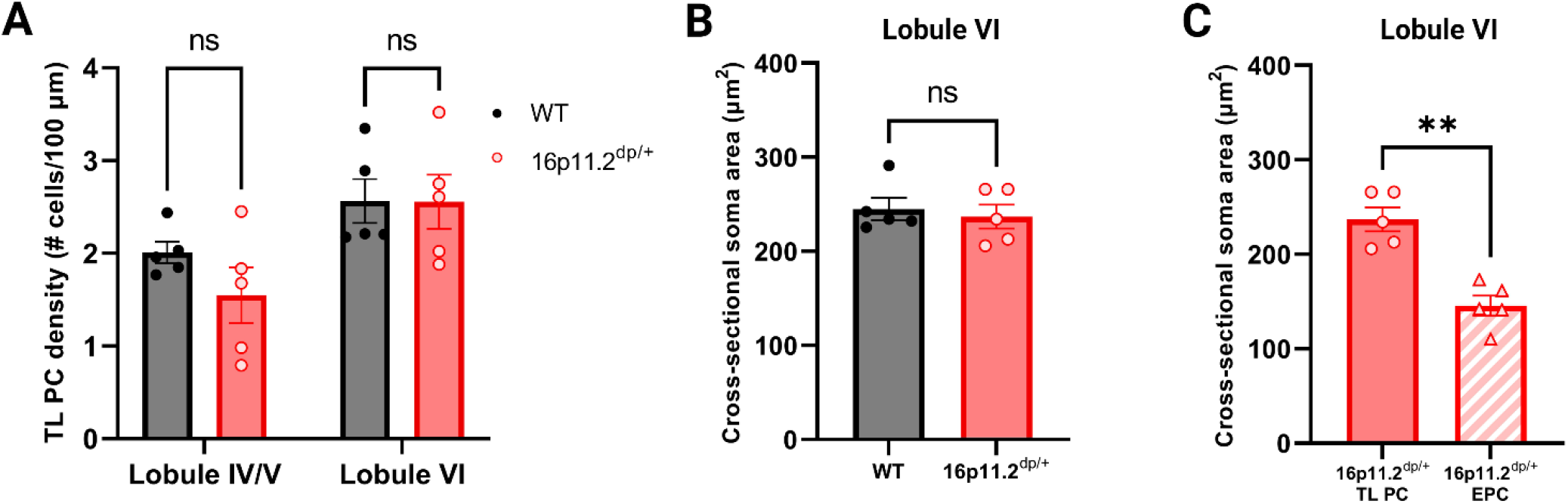
Quantification of density and size of typically located Purkinje cells; size of ectopic Purkinje cells. **A.** No differences were found in typically located Purkinje cell (TL PC) density in lobule IV/V or lobule VI. **B.** No differences were found in typically located Purkinje cell cross-sectional soma area in lobule VI. **A-B:** Mean with SEM, n = 10 (WTs: n = 3 females, n = 2 males; 16p11.2^dp/+^: n = 2 females, n = 3 males), mice were between 3.3-3.5 months old at the time of the experiment. **C.** Typically located Purkinje cells (TL PC) were significantly larger than ectopic Purkinje cells (EPC) (Mean with SEM, n = 5 (16p11.2^dp/+^: n = 2 females, n = 3 males), mice were between 3.3-3.5 months old at the time of the experiment, data from lobule VI.

##### Typically located Purkinje cell somata are significantly larger than ectopic Purkinje cell somata in 16p11.2^dp/+^ mice

In our quantification of ectopic PCs in lobule VI in 16p11.2^dp/+^ mice, we noticed that these cells appeared smaller in size compared to typically located PCs. To quantitatively assess this morphological alteration, we next examined soma cross-sectional area in lobule VI ectopic PCs of 16p11.2^dp/+^ mice relative to typically located PC soma cross-sectional area in 16p11.2^dp/+^ mice. We found a significant decrease in ectopic PC soma cross-sectional area compared to the typically located PC soma cross-sectional area in lobule VI of 16p11.2^dp/+^ mice (unpaired t test, p = 0.0006; 16p11.2^dp/+^ typically located PC mean = 236.8 µm^2^, 16p11.2^dp/+^ ectopic PC mean = 145.6 µm^2^) (**Fig. 2C**). However, it should be noted that primary neurites of ectopic PCs were not always observed and the reduced arborization of ectopic PC dendrites prevented the ability to distinguish between dendrite and axon (**Fig. 1E and 1F**). These two factors could limit our ability to quantify maximal ectopic PC cross-sectional area.

#### Molecular Layer Interneurons

##### Lobule VI-specific decrease in apical parvalbumin+ molecular layer interneurons in 16p11.2^dp/+^ mice

To further explore the effect of the 16p11.2^dp/+^ on the morphology and location of other cell types in the cerebellum, we next compared MLI counts between WT and 16p11.2^dp/+^ mice. We examined overall MLI counts in both lobule IV/V and lobule VI of the cerebellum. In lobule IV/V, no differences were found in DAPI+ cell counts or parvalbumin+ MLI counts (multiple unpaired t tests, Holm – Sidak method, DAPI+ cell counts: p = 0.5308, parvalbumin+ cell counts: p = 0.1697) (**Fig. 3A**). Conversely, in lobule VI, we observed a significant reduction in parvalbumin+ MLIs in 16p11.2^dp/+^ mice compared to WT littermates (multiple unpaired t tests, Holm – Sidak method, parvalbumin+ cell counts: p = 0.0006). However, there was no difference in total cell number in the molecular layer, as measured by DAPI, between 16p11.2^dp/+^ mice and WT littermates in lobule VI (multiple unpaired t tests, Holm – Sidak method, DAPI+ cell counts: p = 0.6172). (**Fig 3B**).

**Figure 3.**
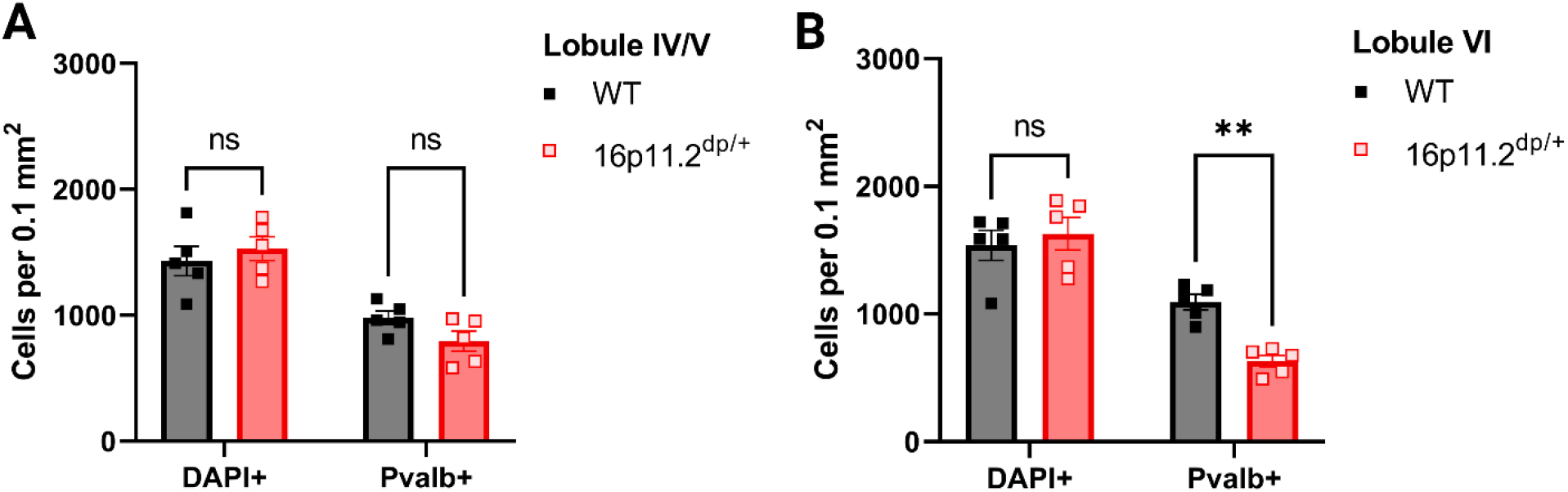
Lobule VI-specific decrease in overall parvalbumin+ molecular layer interneuron cell counts without a decrease in total molecular layer cell density in 16p11.2^dp/+^ mice. **A**. No differences were found in DAPI+ cell counts nor parvalbumin+ (pvalb+) cell counts in lobule IV/V. **B.** In lobule VI, parvalbumin+ (pvalb+) cell counts were significantly decreased in 16p11.2^dp/+^ mice compared to WT littermates, but DAPI+ cell counts were unchanged. **A-B:** Mean with SEM, n = 10 (WTs: n = 3 females, n = 2 males; 16p11.2^dp/+^: n = 2 females, n = 3 males), mice were between 3.3-3.5 months old at the time of the experiment.

We then sought to confirm whether specific subsets of MLIs were reduced in lobule VI of 16p11.2^dp/+^ mice. Early born MLIs reside largely in the basal 1/3 of the molecular layer, while later-born MLIs mostly occupy the apical 2/3 of the molecular layer [65]. When we divided the molecular layer into these two segments (basal 1/3 vs apical 2/3) (**Fig. 4A**), we observed a significant reduction in parvalbumin+ apical 2/3 MLI cell counts in 16p11.2^dp/+^ mice compared to WT littermates (multiple unpaired t tests, Holm – Sidak method, p = 0.005) (**Fig. 4B**). We observed no difference in parvalbumin+ basal 1/3 MLI cell counts (p = 0.9219), nor in overall cell number in the basal 1/3 or apical 2/3 of the molecular layer by DAPI labeling (multiple unpaired t tests, Holm – Sidak method, DAPI+ basal 1/3 cell counts: p = 0.9219, DAPI+ apical 2/3 cell counts: p = 0.9125) (**Fig. 4B**).

**Figure 4.**
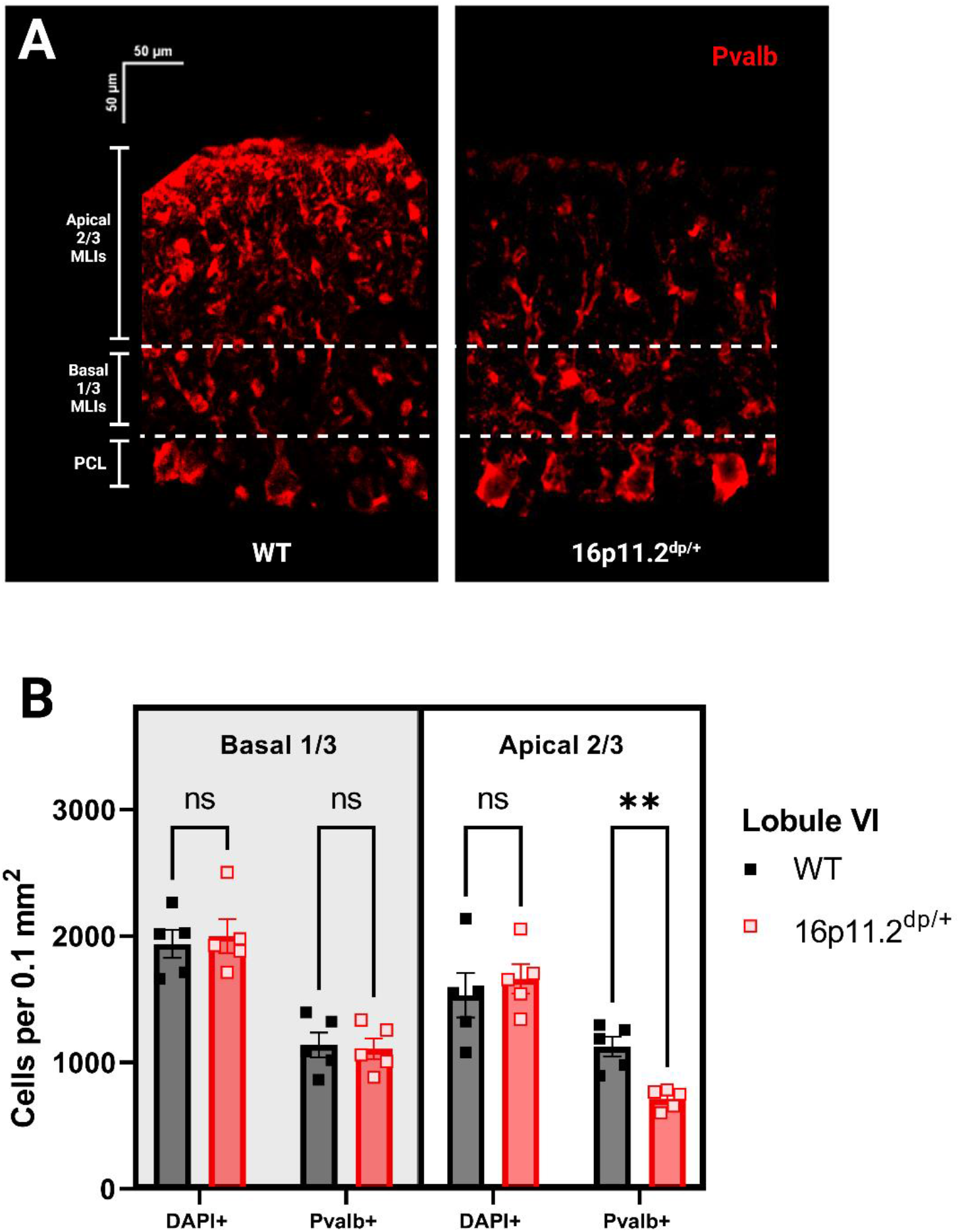
Lobule VI-specific decrease in parvalbumin+ apical MLIs without a decrease in cell density in 16p11.2^dp/+^ mice. **A**. Representative 20X epifluorescent image of parvalbumin (pvalb), shown in red, immunostaining of WT and 16p11.2^dp/+^ tissue in the molecular layer of lobule VI. The molecular layer was divided, as indicated with dashed white lines, into apical 2/3 MLI and basal 1/3 MLI segments and the Purkinje cell layer is indicated as PCL. **B.** In lobule VI, apical 2/3 parvalbumin+ (pvalb+) cell counts were significantly decreased in 16p11.2^dp/+^ mice compared to WT littermates, but apical 2/3 DAPI+, basal 1/3 DAPI+, and basal 1/3 parvalbumin+ cell counts were unchanged. Mean with SEM, n = 10 (WTs: n = 3 females, n = 2 males; 16p11.2^dp/+^: n = 2 females, n = 3 males), mice were between 3.3-3.5 months old at the time of the experiment.

### Behavioral Examination

#### Body Weights

##### No differences in body weight in 16p11.2^dp/+^ mice

We then examined whether 16p11.2 microduplication was associated with changes in body weight, as underweight status has been described as a sex-dependent phenotype in human males with this CNV [68]. We observed a main effect of sex such that male mice were larger than female mice (2-way ANOVA, p = 0.0198). However, there were no significant differences in bodyweight between 16p11.2^dp/+^ mice and WT littermates (2-way ANOVA, no main effect of genotype, p = 0.9736; no genotype by sex interaction, p = 0.9398). Graphical data not shown.

#### Erasmus Ladder

##### No differences in baseline gait or motor coordination in 16p11.2^dp/+^ mice

We investigated the effect of 16p11.2 microduplication on gait using the Erasmus Ladder task. Mice were assessed on days 1-4 for gait abnormalities. Motor coordination is assessed by examining the percentage of incorrect steps (lower rugs) the animal uses to traverse the ladder, defined as missteps. Female mice displayed fewer missteps than male mice in days 1-4 (Type III ANOVA Satterthwaite’s method, main effect of sex, F_1,29_ = 7.1035, p = 0.0125), but there was no main effect of genotype (Type III ANOVA Satterthwaite’s method, no main effect of genotype, F_1,29_ = 3.6905, p = 0.0650) nor interaction effect (Type III ANOVA Satterthwaite’s method, no genotype by sex interaction effect, F_1,29_ = 0.7642, p = 0.3892). However, all groups had fewer missteps as the training progressed (Type III ANOVA Satterthwaite’s method, main effect of day, F_3,86_ = 15.7412, p = 0.0001; day 1 vs. 2 (p = 0.0001), day 1 vs. 3 (p = 0.0001), day 1 vs. 4 (p = 0.0001), day 2 vs. 3 (p = 0.9914), day 2 vs. 4 (p = 0.2205), day 3 vs. 4 (p = 0.3632)) (**Fig. 5A**).

**Figure 5.**
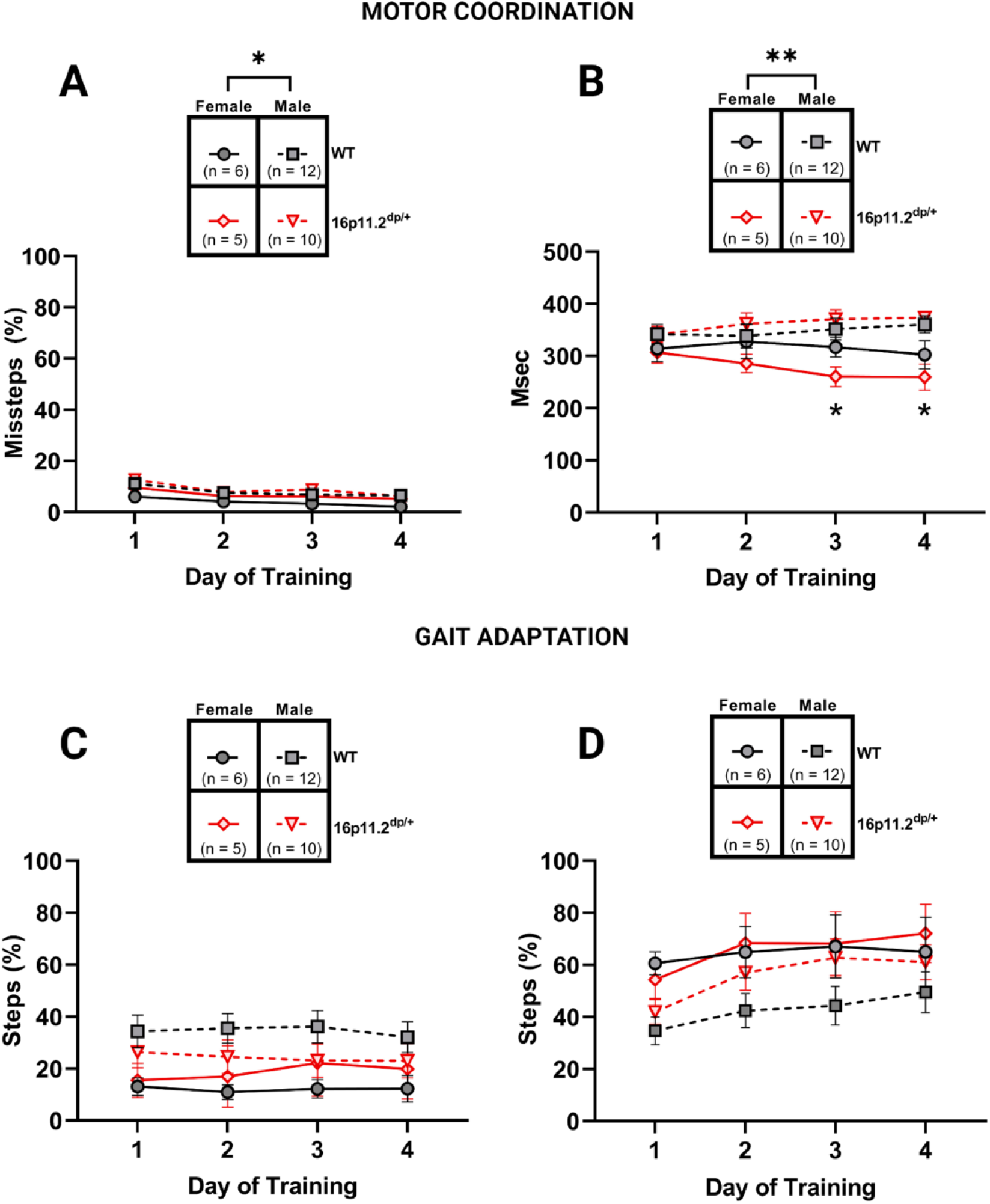
Typical motor coordination and gait adaptation in 16p11.2^dp/+^ mice. **A.** No differences were found in percentage of missteps used between WT and 16p11.2^dp/+^ littermates. However, female mice displayed fewer missteps than male mice. **B.** No differences were found in step time between WT and 16p11.2^dp/+^ littermates. However, female mice were faster than male mice, and their step time increased more than male mice on days 3 and 4 (indicated on graph with asterisk). **C.** No differences were found in the percentage of short steps used. **D.** All mice increased their percentage of long steps used as days of training progressed (main effect of day not indicated on graph). However, there were no sex or genotype differences. **A-D**: Mean with SEM, n = 33 (WTs: n = 6 females, n = 12 males; 16p11.2^dp/+^: n = 5 females, n = 10 males), mice were between 3.3-3.5 months old at the time of the experiment.

We also explored average step times to determine if 16p11.2 microduplication altered step speed. Male mice displayed significantly slower step times than female mice (Type III ANOVA Satterthwaite’s method, main effect of sex, F _1,29_ = 9.5058, p = 0.005). Additionally, female mice displayed significantly faster step times as the days of training progressed (Type III ANOVA Satterthwaite’s method, main effect of sex, F _1,29_ = 9.5058, p = 0.0045; with a sex by day interaction, F _3,86_ = 3.6530, overall p = 0.016, day 1 (p = 0.1718), day 2 (p = 0.0565), day 3 (p = 0.0017), day 4 (p = 0.003); no main effect of genotype, F _1,29_ = 0.417, p = 0.5235)) (**Fig. 5B**). These data confirm that all groups display typical motor coordination.

As mice learn the Erasmus Ladder task, they transition from using short steps (stepping on each rung) to long steps (stepping on every other rung) because long steps are a more efficient means of crossing the ladder. The animal’s ability to transition from short to long steps is a measure of gait adaptation. We observed no interaction effects nor main effects of genotype or sex for short steps (Type III ANOVA with Satterthwaite’s method, no main effect of genotype, F _1,29_ = 0.0541, p = 0.8178; or sex, F _1,29_ = 3.7623, p = 0.0622) on days 1-4 (**Fig. 5C**). There were no interaction effects nor main effects of genotype or sex for long steps on days 1-4 (Type III ANOVA with Satterthwaite’s method, no main effect of genotype, F _1,29_ = 0.7058, p = 0.4077; or sex, F _1,29_ = 3.668, p = 0.0654), however, all groups significantly increased their percentage of long steps used as the days of training progressed (Type III ANOVA with Satterthwaite’s method, main effect of day, F_3,86_ = 17.8853, p = 0.0001; day 1 vs. 2 (p = 0.0001), day 1 vs. 3 (p = 0.0001), day 1 vs. 4 (p = 0.0001), day 2 vs. 3 (p = 0.5198), day 2 vs. 4 (p = 0.3166), day 3 vs. 4 (p = 0.9857)) (**Fig. 5D**). These data confirm that all groups display typical gait adaptation.

##### Cerebellar-dependent associative motor learning differences in female 16p11.2^dp/+^ mice

Although 16p11.2 microduplication does not appear to affect gait adaptation and motor coordination, we hypothesized that it may affect cerebellar associative motor learning. In addition to assessing gait abnormalities, the Erasmus Ladder task also examines tone-cued, cerebellar-dependent associative motor learning on days 5-8. This form of cerebellar-dependent learning is accomplished by presenting a tone prior to raising a rung in the path of a traversing animal, in which the animal learns to use the tone cue to time their jump over the raised ladder rung to avoid the obstacle [45]. We observed an interaction between genotype and sex in the pre-perturbation step of days 5-8, driven mostly by a significantly faster step time in 16p11.2^dp/+^ females compared to WT females and 16p11.2^dp/+^ males (Type III ANOVA Satterthwaite’s method, genotype by sex interaction, F _1,29_ = 8.4867, p = 0.007; with WT vs. 16p11.2^dp/+^ males (p = 0.0730), WT vs. 16p11.2^dp/+^ females (p = 0.0326), WT male vs. female (p = 0.0561), and 16p11.2^dp/+^ male vs. Female (p = 0.0425); no main effect of genotype, F _1,29_ = 0.5378, p = 0.4694; no main effect of sex, F _1,29_ = 0.0417, p = 0.8396). There was also a main effect of day, such that all groups’ step times became significantly faster as the days of training progressed (Type III ANOVA with Satterthwaite’s method, main effect of day, F_3,86_ = 22.0529, p = 0.007; day 1 vs. 2 (p = 0.0001), day 1 vs. 3 (p = 0.0001), day 1 vs. 4 (p = 0.0001), day 2 vs. 3 (p = 0.7025), day 2 vs. 4 (p = 0.9912), day 3 vs. 4 (p = 0.8599)) (**Fig. 6A**). Additionally, we observed an interaction between genotype and sex in the post-perturbation step, also driven by a significantly faster step time in 16p11.2^dp/+^ females compared to WT females in days 5-8 ((Type III ANOVA Satterthwaite’s method, genotype by sex interaction, F _1,29_ = 22.0529, p = 0.0098) with WT vs. 16p11.2^dp/+^ males (p = 0.2589) and WT vs. 16p11.2^dp/+^ females (p = 0.0151); no main effect of genotype, F _1,29_ = 2.0288, p = 0.1654; no main effect of sex, F _1,29_ = 0.0243, p = 0.8773). There was also a main effect of day, such that all groups became significantly faster as the days of training progressed (Type III ANOVA with Satterthwaite’s method, main effect of day, F_3,86_ = 22.0529, p = 0.0001; day 1 vs. 2 (p = 0.0001), day 1 vs. 3 (p = 0.0001), day 1 vs. 4 (p = 0.0001), day 2 vs. 3 (p = 0.7137), day 2 vs. 4 (p = 0.9982), day 3 vs. 4 (p = 0.8100)) (**Fig. 6B**). These data indicate tone-cued cerebellar associative motor learning occurred for all groups, but the 16p11.2^dp/+^ females show a significantly faster step time relative to WT females and 16p11.2^dp/+^ males.

**Figure 6.**
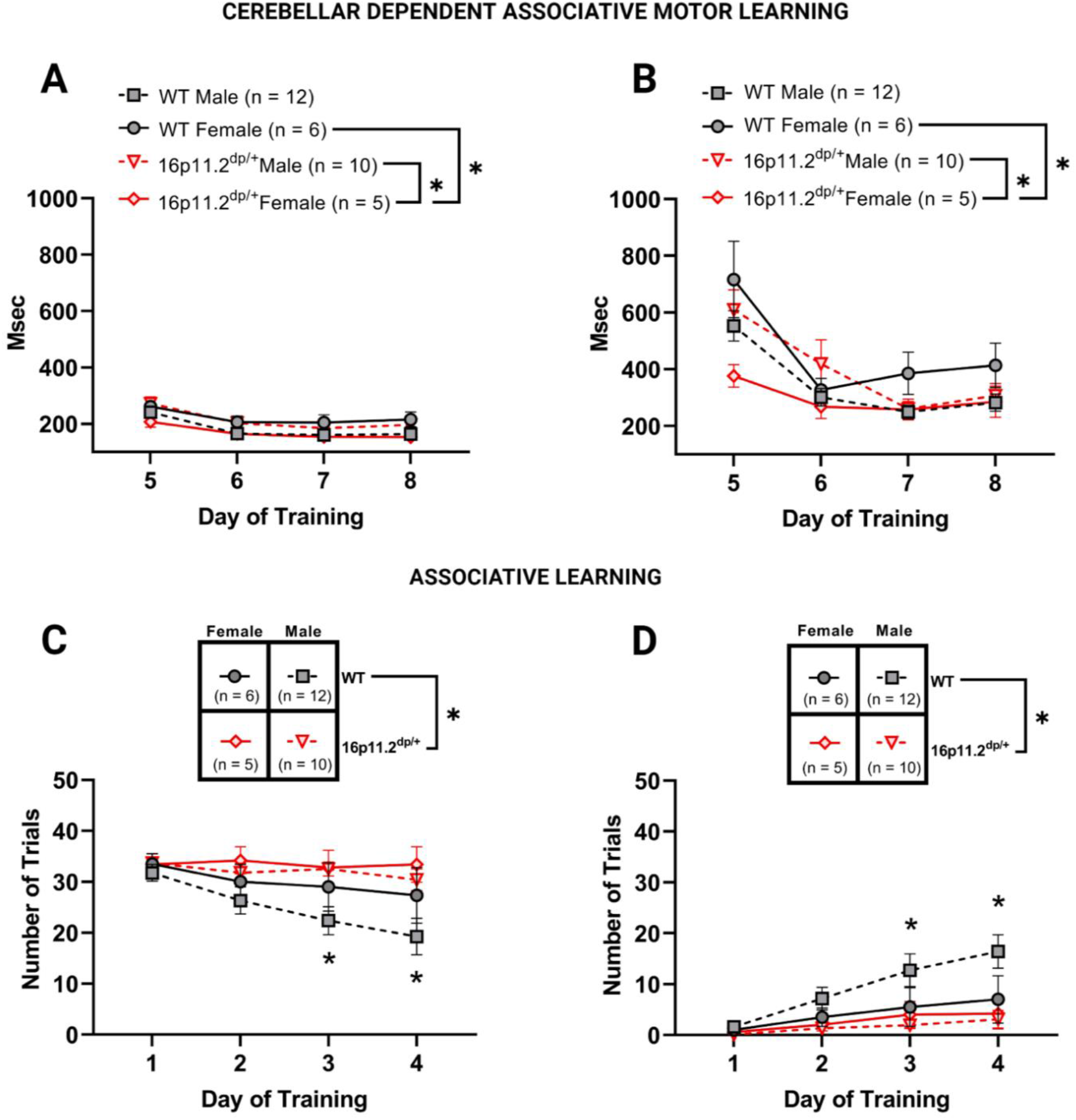
Cerebellar-dependent learning differences in female 16p11.2^dp/+^ mice; associative learning impairments in 16p11.2^dp/+^ mice. **A.** Female 16p11.2^dp/+^ mice were significantly faster on the steps immediately preceding the perturbation than 16p11.2^dp^/^+^ males and WT females. **B.** Female 16p11.2^dp/+^ mice were significantly faster on the steps immediately following the perturbation than 16p11.2^dp^/^+^ males and WT females. **C.** Overall, 16p11.2^dp/+^ mice left on air cue more frequently than WT littermates. This difference was most evident on days 3 and 4 of training (indicated on graph with asterisk). **D.** Overall, WT mice left on light cue more frequently than 16p11.2^dp/+^ mice. This difference was most evident on days 3 and 4 of training (indicated on graph with asterisk). **A-D**: Mean with SEM, n = 33 (WTs: n = 6 females, n = 12 males; 16p11.2^dp/+^: n = 5 females, n = 10 males), mice were between 3.3-3.5 months old at the time of the experiment.

##### Associative learning impairments in 16p11.2^dp/+^ mice

Mice are also assessed on Erasmus Ladder days 1-4 for associative learning of trial cues, which was measured by the ability to learn visual (a light) and sensory (a tailwind of pressurized air) start cues for each trial. For days 1-2, we found most animals did not respond to the light cue, with no differences between genotypes, but the groups began to separate by genotype in days 3 and 4. WT mice learned to associate the light cue with the initiation of a trial on days 3 and 4 while 16p11.2^dp/+^ mice did not (Type III ANOVA Satterthwaite’s method, main effect of genotype, F _1,29_ = 4.9755, p = 0.0336; genotype by day interaction, F_3,86_ = 3.7700, p = 0.0136; day 1 (p = 0.7078), day 2 (p = 0.1462), day 3 (p = 0.0201), day 4 (p = 0.0021); no main effect of sex, F _1,29_ = 0.9314, p = 0.3425) (**Fig. 6D**). Moreover, as mice learn the Erasmus Ladder task, they use the light cue more frequently to avoid the unpleasant air cue. We observed that 16p11.2^dp/+^ mice left on air cue more frequently than WT mice on day 3 and 4 of training (Type III ANOVA Satterthwaite’s method, main effect of genotype, F _1,29_ = 4.6359, p = 0.0398; with a genotype by day interaction, F_3,86_ = 2.6502, p = 0.0500; day 1 (p = 0.7592), day 2 (p = 0.1129), day 3 (0.0298), day 4 (p = 0.0057); no main effect of sex, F _1,29_ = 1.6332, p = 0.2114) (**Fig. 6C**). These data suggest that because 16p11.2^dp/+^ mice are leaving on the air cue more frequently than WTs, 16p11.2^dp/+^ mice are failing to associate the light cue with the trial start. 16p11.2^dp/+^ mice may also be less inclined to leave the goalbox. Additionally, these data suggest WT mice learn better than 16p11.2^dp/+^ mice to associate the light cue with the start of a trial to avoid the air cue.

#### Eyeblink conditioning

##### Cerebellar-dependent associative learning impairments in 16p11.2^dp/+^ mice

Acquisition of delay eyeblink conditioning with a 400 ms interstimulus interval was tested between WT and 16p11.2^dp/+^ mice. Delay eyeblink conditioning with a relatively long interstimulus interval was used to assess potential deficits in cerebellar learning and timing in 16p11.2^dp/+^ mice that also showed abnormalities in PC organization and reduction in apical 2/3 MLI parvalbumin expression. Repeated measures ANOVA on the CR percentage data for WT and 16p11.2^dp/+^ mice across acquisition of delay eyeblink conditioning revealed a significant interaction of genotype and session factors (F_(9,117)_ = 3.692, p = 0.001). There was also a significant main effect of session (F_(9, 117)_ = 26.086, p = 0.001). Post-hoc tests (Fisher’s Least Significant Difference (LSD), p = 0.016) revealed that WT mice showed more CRs on session 10 relative to 16p11.2^dp/+^ mice (**Fig. 7A**). Both groups showed a progressive increase in CR percentage across sessions (for all comparisons, p = 0.045). Furthermore, we observed no significant interaction between CR percentage and sex (F_(9,117)_ = 1.542, p = 0.4562). Although 16p11.2^dp/+^ mice showed a larger CR percentage earlier in training, they failed to reach the same level of learning as the WT mice by the final session. This result provides evidence that WT mice show a higher level of cerebellar dependent learning compared to 16p11.2^dp/+^ mice.

**Figure 7.**
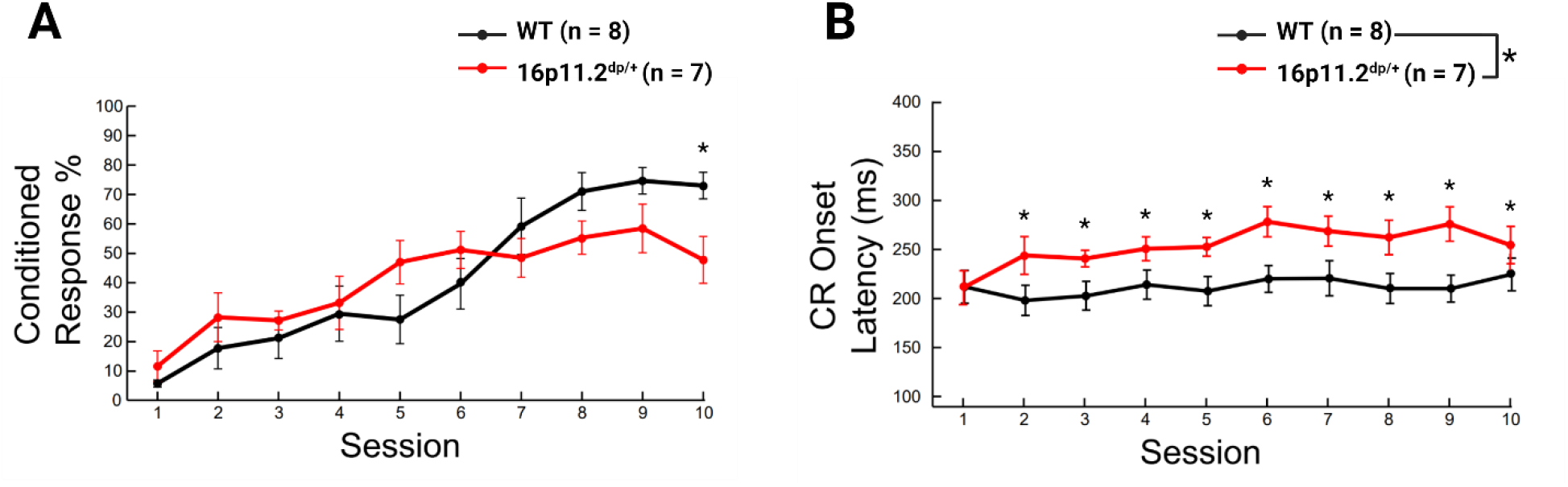
Impaired acquisition of conditioned response and increased conditioned response onset latencies in 16p11.2^dp/+^ mice. **A.** 16p11.2^dp/+^ mice showed fewer conditioned responses on session 10 of training relative to WT mice (indicated on graph with asterisk). **B.** The onset of the conditioned response (CR) in 16p11.2^dp/+^ mice was significantly later than that of WT littermates (indicated on graph with asterisk). **A-B**: Mean with SEM, n = 15 (WTs: n = 3 females, n = 8 males; 16p11.2^dp/+^: n = 5 females, n = 2 males), mice were between 4.8-6.8 months old at the time of the experiment.

##### Increased conditioned response onset latency in 16p11.2^dp/+^ mice

The cerebellum controls the adaptive timing of CR expression during eyeblink conditioning [32, 34, 35, 69]. The mechanism controlling this adaptive timing is learned decreases in simple spike activity of specific PCs that receive complex spikes elicited by the US (air puff). The variability in CR onset time increases as a function of interstimulus duration, which is also accompanied by an increase in individual PC variability with respect to decreases in simple spikes before CR onset [32–34]. 16p11.2^dp/+^ mice with abnormalities in the region of the cerebellar cortex necessary for eyeblink conditioning (lobule VI) [35, 36] would thus be expected to show deficits in CR timing. Repeated measures ANOVA on the CR onset data for WT and 16p11.2^dp/+^ mice across acquisition of delay eyeblink conditioning revealed significant main effects of genotype (F_(1,13)_ = 5.024, p = 0.05) and session (F_(9,117)_ = 2.508, p = 0.05) factors. Post hoc tests (LSD, p = 0.043) revealed that 16p11.2^dp/+^ mice showed significantly later CR onsets relative to WT mice from session 2 though session 10 **(Fig. 7B**). Both groups of mice showed improved CR timing over the course of training as CR onset time progressively approached the timing of the US. By the end of training, the improvement in CR onset timing from session 2 to session 10 was less robust in 16p11.2^dp/+^ mice compared with the improvement observed in WT mice. These results suggest that 16p11.2^dp/+^ mice are impaired in expression and learned improvement of adaptive timing during cerebellar driven behavior. 16p11.2^dp/+^ mice not only showed fewer CRs over the course of training, but when they showed CRs, the onset latency was consistently impaired.

#### Prepulse Inhibition

##### Impaired sensorimotor gating in female 16p11.2^dp/+^ mice

The 16p11.2 microduplication is linked to neuropsychiatric disorders with reduced sensorimotor gating such as SCZ and ASD [60–62]. Reduction in PPI is believed to be linked to dysfunction in the sensorimotor gating mechanism [70]. To test sensorimotor gating and startle response, we used the PPI task. We observed an expected main effect of decibel, such that higher decibel prepulses elicited greater PPI (Type III ANOVA Satterthwaite’s method, main effect of decibel, F_2,32_ = 19.7922, p = 0.0001). There was also a small decibel by genotype interaction effect (Type III ANOVA Satterthwaite’s method, decibel by genotype interaction effect, F_2,32_ = 3.2950, p = 0.05; WT 10 dB vs. 15 dB (p = 0.0085); 16p11.2^dp/+^ mice 10 dB vs. 15 dB (p = 0.0286)). We did not observe consistent PPI response percentages above 0 for 5 dB in WT mice, but we did observe PPI in WT mice at 10 dB and 15 dB. There were no main effects of genotype or sex (Type III ANOVA Satterthwaite’s method, no main effect of genotype, F_1,16_ = 0.1696, p = 0.6859; no main effect of sex, F_1,16_ = 0.1119, p = 0.7423). Interestingly, we observed a genotype by sex interaction effect (Type III ANOVA Satterthwaite’s method, genotype by sex interaction effect, F_1,16_ = 9.3566, p = 0.0075). Specifically, follow up tests indicated that female 16p11.2^dp/+^ mice display reduced PPI response percentage compared to WT females and 16p11.2^dp/+^ males (WT females vs. 16p11.2^dp/+^ females (p = 0.0144), WT males vs. 16p11.2^dp/+^ males (p = 0.1068), WT females vs. WT males (p = 0.0720), 16p11.2^dp/+^ females vs. 16p11.2^dp/+^ males (p = 0.0289)) **(Fig. 8A**). Therefore, these data show sex-specific deficits in sensorimotor gating in female 16p11.2^dp/+^ mice relative to WT females and 16p11.2^dp/+^ males.

**Figure 8.**
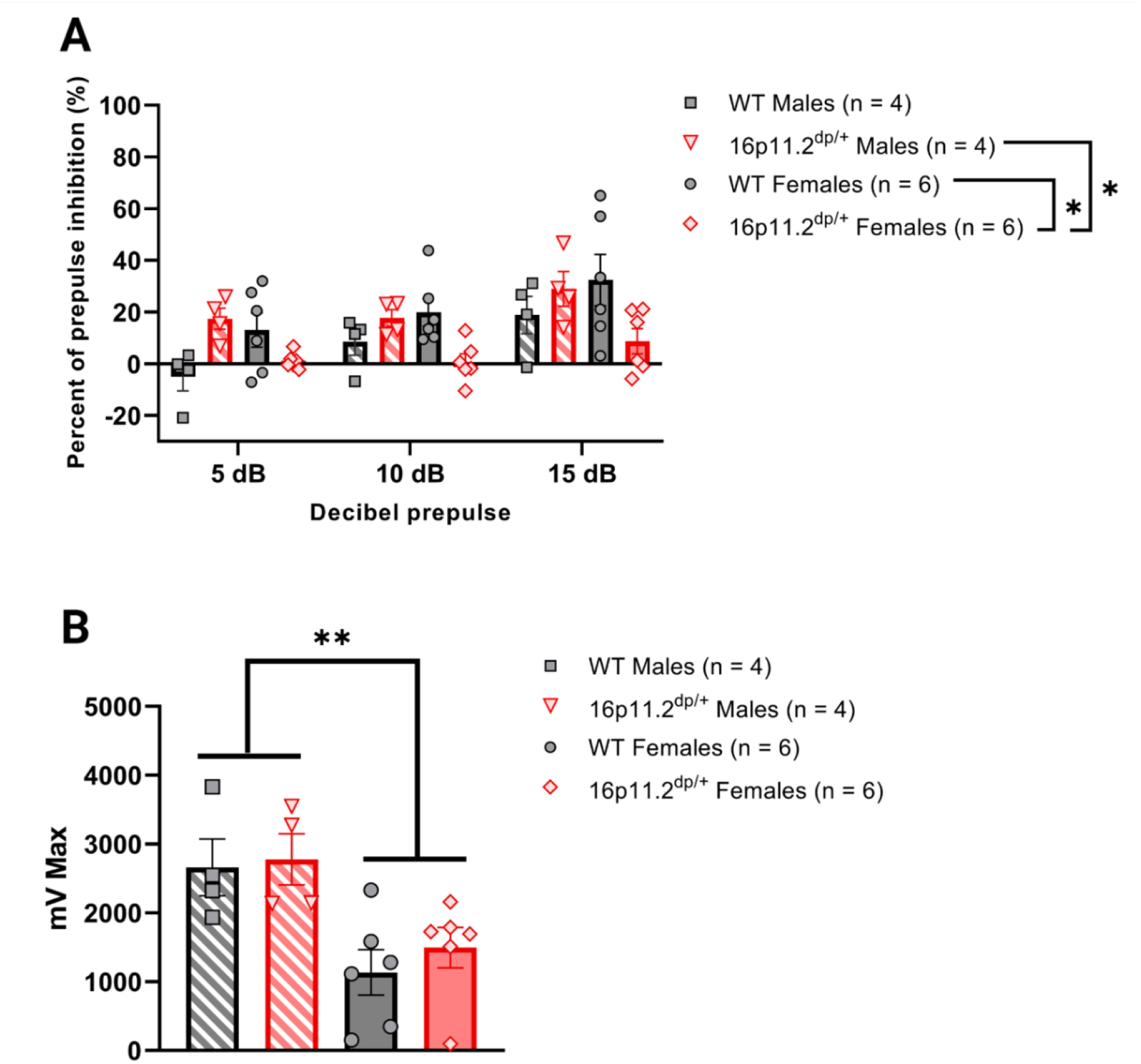
Impaired sensorimotor gating in female 16p11.2^dp/+^ mice; greater startle response in male mice. **A.** Female 16p11.2^dp/+^ showed reduced prepulse inhibition relative to WT females and 16p11.2^dp/+^ males. **B.** Male mice display a greater startle response than female mice (Shown: block II, not shown: block I and III). **A-B**: Mean with SEM, n = 20 (WTs: n = 6 females, n = 4 males; 16p11.2^dp/+^: n = 6 females, n = 4 males). All mice were between the ages of 2.7 – 4.8 months old at the time of the experiment.

##### Increased startle response in male mice

For startle response, we did not observe a genotype or interaction effect (Type III ANOVA Satterthwaite’s method, no main effect of genotype, F_1,16_ = 0.5108, p = 0.4811). There was habituation to startle response across the duration of the experiment for all groups, but the startle response was detectable throughout the experiment (Type III ANOVA Satterthwaite’s method, main effect of block, F_2,32_ = 15.2470, p = 0.0001). We observed a main effect of sex, such that male mice displayed significantly higher responses to the auditory startle pulse compared to female mice (Type III ANOVA Satterthwaite’s method, main effect of sex, F_1,16_ = 12.2470, p = 0.0026). These data indicate sex differences are present in response to the auditory startle pulse (**Fig. 8B**).

## DISCUSSION

### Summary

Previous studies of 16p11.2^dp/+^ mice have largely focused on cortical abnormalities, despite the growing evidence pointing to cerebellar alterations contributing to neuropsychiatric phenotypes. Here, we took a histological and behavioral approach to investigate the effects of this CNV in the mouse cerebellum. We have identified novel alterations to cerebellar structure and cerebellar-dependent behavior in 16p11.2^dp/+^ mice. Our histological data indicate that this CNV leads to lobule VI-specific alterations in PC localization and expression of parvalbumin in MLIs, specifically in the apical 2/3 of the molecular layer. Furthermore, our behavioral data suggests that the 16p11.2^dp/+^ CNV does not impair gait or motor coordination, as measured by the Erasmus Ladder, but it does impair cerebellar-dependent learning as measured by delay eyeblink conditioning. In addition, other behavioral assessments, such as PPI and the associative learning aspects of the Erasmus Ladder task, reveal that 16p11.2^dp/+^ mice display similar phenotypes to human patients with this CNV with respect to deficits in sensorimotor gating [60–62], associative learning [71], and cerebellar dependent associative learning [37–42].

### Possible cellular mechanisms underlying delay eyeblink conditioning deficits in 16p11.2^dp/+^ mice

Our histological and behavioral analyses of the 16p11.2^dp/+^ mouse cerebellum revealed structural alterations in lobule VI together with deficits in cerebellar dependent associative motor learning assessed through delay eyeblink conditioning, a behavior that is associated with lobule VI [35, 36]. Lobule-specific phenotypes in 16p11.2^dp/+^ mice become more apparent when considering that we did not find any structural differences in lobule IV/V, nor did we find differences in the motor behaviors associated with this lobule, such as gait adaptation and motor coordination [26, 47]. Therefore, our results suggest a possible mechanistic link between lobule VI structural alterations and the behavioral deficits we have observed in 16p11.2^dp/+^ mice.

In delay eyeblink conditioning, 16p11.2^dp/+^ mice exhibited longer CR onset latencies combined with impaired CR acquisition. This behavioral phenotype suggests a lack of variability in PC learning-related firing changes, which control CR kinematics [32–34], and could be a result of the lobule VI-specific structural alterations identified in 16p11.2^dp/+^ mice. While we do not know exactly what role ectopic PCs or reduced parvalbumin expression in apical 2/3 MLIs play in the underlying eyeblink circuitry in lobule VI, we posit that alterations to either or both cell types could result in an imbalance of simple and complex spike output of typically located PCs onto cerebellar nuclei.

Specifically, based on the granule layer localization of ectopic PCs in lobule VI in 16p11.2^dp/+^ mice, ectopic PC dendrites may synapse onto granule cell axons (parallel fibers), or even receive direct input from mossy and climbing fibers, leading to aberrant inputs onto ectopic PCs. Furthermore, PC axons project to the cerebellar nuclei before they migrate through the intermediate zone to establish the PC monolayer during development [72], indicating that ectopic PC axons may be capable of providing signaling output to cerebellar nuclei despite their location in the granule layer. If ectopic PCs exhibit the same spike activity of typically located PCs, the possible inputs of parallel fibers (input that can also produce simple spiking in PCs, aside from tonic PC firing) and climbing fibers (the input responsible for complex spiking in PCs) [73] onto ectopic PCs could lead to the summation of atypical simple and complex spike output from lobule VI to the cerebellar nuclei. Although ectopic PCs are found in various cerebellar lobules in both WT and 16p11.2^dp/+^ mice, the notable increase in this cell type in lobule VI of 16p11.2^dp/+^ mice may disrupt the signaling in lobule VI beyond a point at which typical behavior can be produced. Moreover, MLIs are believed to modulate the activity of typically located PCs [74]. If MLI inhibitory neurotransmission is decreased due to decreased parvalbumin expression in apical 2/3 MLIs, this could also alter the balance and regularity of complex and simple spike output from typically located PCs. In other mouse models, loss of apical 2/3 MLI GABAergic neurotransmission increased the regularity of PC simple spike firing and increased the complex spike firing rate [65]. Lack of PC firing variability in 16p11.2^dp/+^ mice may be due to reduced parvalbumin expression in apical 2/3 MLIs, leading to a decrease of inhibitory inputs onto typically located PCs, thus, altering the timing of typically located PC activity through simple spikes.

In addition, an increase in complex spike firing rate, as described from loss of apical 2/3 MLI GABAergic neurotransmission [65], may result in the learning impairments that we observed in 16p11.2^dp/+^ mice during eyeblink conditioning. For example, if complex spikes are firing more often, typically located PCs could be firing complex spikes when there is no learning input, and therefore, impair learning by weakening the association. In other mouse models, lobule VI-specific MLI inactivation (∼20%) via DREADDs has resulted in similar impaired delay eyeblink conditioning CRs [75]. Together, these data suggest that the learning deficits we observed in 16p11.2^dp/+^ mice may be a result of the cerebellar abnormalities observed in the distribution of PCs and/or apical 2/3 MLI parvalbumin expression in lobule VI. However, further work is needed to assess the electrophysiological activity of ectopic PCs, typically located PCs, and apical MLIs during eyeblink conditioning to better understand the link between the structural and behavioral alterations we have identified in 16p11.2^dp/+^ mice.

### Ectopic Purkinje cell phenotypes in other mouse models

The occurrence of ectopic PCs in the granule layer with preserved cerebellar foliation and typically located PC density is reported rarely in other mutant mouse models, as is the morphological (soma area) descriptions of ectopic PCs. However, one mutant mouse model that displays a comparable ectopic PC phenotype to 16p11.2^dp/+^ mice, is the p35 knockout (KO), in which mice lack the neuronal-specific activator (p35) of cyclin-dependent kinase 5 (Cdk5). Cdk5, along with its regulatory subunit p35, is necessary for neuronal migration and laminar formation [76]. This mouse model also displayed ectopic PCs in the granule layer without gross alterations to cerebellar foliation or typically located PC density, although the lobule these ectopic PCs were observed in was not specified [76].

In contrast to the ectopic PCs found in 16p11.2^dp/+^ mice and the p35 KO model [76], other examples of mutant mouse models with ectopic PCs display vastly reduced or no cerebellar foliation, ectopic PC clusters, lack of a PC monolayer, and aberrant lamina. These mutant mouse models include: *Reeler* [77–79], *Scrambler* [80, 81], *VLDLR/ApoE2* [82], *Src/Fyn* [83], *Straggerer* [84], *Pten* [85], *SmoA2* [86, 87], and *Cxcr4-/SDF-1* deficient [88–90]. While each of these mutations targets different genes, the commonality between each model is the disruption of PC migration. This disruption can result from numerous factors, such as alterations to the Reelin signaling pathway (as seen in *Reeler, Scrambler, VLDLR/ApoE2, and Src/Fyn* mutants), SHH signaling (in *SmoA2* and *Stragger* mutants), or Bergmann glia gene expression (as observed in *Pten* mutants). While 16p11.2^dp/+^ mice did not exhibit as great a number of ectopic PCs as the mouse models described above, the mRNA expression of two of the genes in these models, *Dab1* (*Scrambler* mutant) and *RORα* (*Straggerer* mutant, a gene expressed by PCs and MLIs, which is also required for cerebellar development) were found to be decreased by about 10-11% in the cerebellum of 16p11.2^dp/+^ mice [58]. These data indicate that ectopic PC phenotypes vary across a spectrum of characteristics, potentially resulting from alterations to distinct mechanisms. However, it is likely that shared developmental and migratory genes play a role in the formation of ectopic PCs, regardless of the extent to which this cell type is present.

### Commonality and origins of ectopic Purkinje cells

Furthermore, our discovery of ectopic PCs in the granule layer of *both* WT and 16p11.2^dp/+^ mice raises intriguing questions about their commonality and origins. Other groups have also identified small numbers of ectopic PCs present in WT rats and rabbits in a variety of lamina of the cerebellum, including the cerebellar nuclei and the molecular layer, in addition to the granule layer [91]. Another mouse model, phospholipase D2 (PLD2) KO, reported ectopic PCs in the molecular layer and arbor vitae in *both* their WT and PLD2 KO mice, however to a greater extent in their KOs [92]. Additionally, the cerebella of these animals were typical in size and foliation [91, 92].

Regarding the origins of ectopic PCs in both WT and 16p11.2^dp/+^ mice, we suggest that these are cells that fail to migrate to the PC layer of the cerebellum during development. Ectopic PCs may be present in the granule layer of adult WT and 16p11.2^dp/+^ mice as a result of PCs being trapped in the intermediate zone during their migratory path from the ventricular zone to the cerebellar cortical plate between E18-P5 [93]. Based on the previously described findings of ectopic PCs in WT rats, rabbits, and mice [91, 92], it is possible that this is a naturally occurring phenomenon of cerebellar development. Some trapped PCs may degenerate and undergo apoptosis, while others will remain in ectopic locations and continue differentiating, as typically localized PCs begin differentiating on their migratory path to the cerebellar cortical plate [86], albeit to a lesser extent than their typically located counterparts [91]. Given that we were able to label these ectopic PCs with standard PC markers (calbindin and parvalbumin), these cells appear to have continued some aspect of typical PC differentiation despite their aberrant localization, although these ectopic PCs also exhibited reduced soma size. Further analysis is needed to determine if WT ectopic PCs in lobule IV/V/VI and 16p11.2^dp/+^ ectopic PCs in lobule IV/V also exhibit reduced soma area.

It is also important to note that we cannot explain why ectopic PCs were significantly increased in 16p11.2^dp/+^ mice in lobule VI but not lobule IV/V, or at what point their prevalence would influence behavior. It is possible that the deviating frequency of ectopic PCs in WT vs. 16p11.2^dp/+^ mice could be due to differential gene expression associated with this CNV; altering pathways that affect PC migration, the PC progenitor pool, or PC maturation. In support, a study of primary cortical neurons isolated from 16p11.2^dp/+^ mice revealed an abnormal developmental phenotype, posited by the authors to involve premature closure of the critical period and accelerated GABAergic development, driven by *Taok2* over-expression (one of the duplicated genes on the 16p11.2 locus) and subsequent over-activity of JNK [94]. If these findings extend to the cerebellum, it is possible that accelerated GABAergic PC development could contribute to a significant increase of ectopic PCs in 16p11.2^dp/+^ mice. To better understand this phenomenon, further investigation is required to explore the presence of ectopic PCs in other cerebellar lobules in both genotypes. It will be interesting to examine in future studies what timepoint of development ectopic PCs appear in both genotypes to further elucidate their origins.

### No alterations to typically located Purkinje cells in 16p11.2^dp/+^ mice

In contrast to our findings, reduced PC density and PC soma area in the cerebellar vermis and hemispheres has been previously reported in the postmortem brain tissue of individuals with ASD [49–51] and SCZ [53]. Additionally, PC loss has been reported specifically in lobule VI and VII in individuals with ASD [52]. However, others have reported no loss of typically located PCs in individuals with SCZ [54]. In 16p11.2^dp/+^ patient-derived iPSC forebrain cortical neurons, reduced soma size and dendritic length was observed [95], but this finding was not replicated in 16p11.2^dp/+^ mice PC soma cross-sectional area. It is possible our results do not match those described above due to investigation of different brain regions (cerebellum vs. forebrain), different models (mouse vs. human vs. iPSC) or differing etiologies of ASD vs. SCZ vs. 16p11.2 microduplication. Additional exploration of the cerebellum in this CNV is required to ascertain the presence of any other alterations to typically located PC morphology that may still be unidentified, such as changes in dendritic arborization.

### Reduced parvalbumin expression without loss of neuronal density in 16p11.2^dp/+^ mice

Our findings of reduced parvalbumin labeling in apical MLIs in lobule VI without a concomitant reduction in total molecular layer cells (as quantified by DAPI), suggests that 16p11.2^dp/+^ mice display reduced parvalbumin expression in the apical MLIs of lobule VI without an overall loss of MLI density. Recent bulk RNA sequencing using the same 16p11.2 microduplication mouse model in this study supports this hypothesis, having found a ∼10% reduction in parvalbumin mRNA expression in the cerebellum [58]. While mRNA expression does not necessarily reflect protein expression levels, these data further support our hypothesis that the reduction we observed in parvalbumin+ apical 2/3 MLI counts in lobule VI, but not lobule IV/V, is due to a decrease in parvalbumin expression in these neurons, and not a loss of neuronal density. Interestingly, the previously described developmental phenotype observed in a study of primary cortical neurons isolated from 16p11.2^dp/+^ mice also revealed that parvalbumin mRNA expression was decreased in the prefrontal cortex and hippocampus of adult 16p11.2^dp/+^ mice; the authors theorized that parvalbumin expression may be downregulated in adulthood as a compensatory mechanism in attempt to restore typical excitation/inhibition balance from accelerated GABAergic neuronal development [94]. Additional investigation is required to validate the loss of parvalbumin protein expression in cerebellar lobule VI of 16p11.2^dp/+^ mice. Moreover, it is necessary to quantify parvalbumin+ and DAPI+ cell counts in other lobules to determine whether the decrease in parvalbumin+ MLI cell counts is present in other cerebellar lobules.

### Reduced parvalbumin expression in other brain regions in humans and mouse models related to neuropsychiatric disorders

Other studies of SCZ in postmortem human brains, SCZ rat models (methylazoxymethanol acetate (MAM) G17 model), as well as ASD mouse models (*Shank)* have similarly reported a reduction in parvalbumin mRNA expression and/or parvalbumin+ cell counts, but not loss of parvalbumin+ neuron density. In the *Shank* ASD mouse model, both parvalbumin+ cell number (as quantified via immunohistochemistry) and parvalbumin protein expression (as quantified via western blot) were reduced in the somatosensory cortex and striatum, but total neuron density was unchanged [96]. MAM-treated rats displayed a significantly decreased density of parvalbumin+ interneurons throughout the medial prefrontal cortex and ventral subiculum of the hippocampus, however, total cell counts were not reported [97].

In postmortem prefrontal cortex samples of individuals with SCZ, *in situ* hybridization and trans-illuminated autoradiographic film images revealed reduced parvalbumin mRNA expression and no change in total neuron number. Interestingly, this reduction in parvalbumin mRNA expression was layer-specific, such that layer III and IV of the PFC in individuals with SCZ exhibited significantly reduced parvalbumin mRNA expression while layers I, II, and V/VI did not [98]. Cortical layer development, similarly to MLI development, proceeds in an inside-out fashion, such that later born neurons migrate past early born neurons [65, 99]. Although we observed reduced parvalbumin+ cell density in the apical 2/3 of the molecular layer in lobule VI of the 16p11.2^dp/+^ mouse cerebellum, it is fascinating that the reduction of parvalbumin mRNA expression has been reported in later-born layers in other areas of the brain in individuals with SCZ as well.

### Lobule specificity and cerebellar functional subregions in humans and mice

Our histological analyses point to a lobule VI-specific phenotype and thus leads into the examination of the functional subregions within the cerebellum. Many mouse and human lesion studies, as well as fMRI studies in humans, point to functional gradients within the cerebellum, such that motor tasks are represented in anterior cerebellar lobules (lobule I – anterior lobule VI), with a second sensorimotor representation in lobule VIII, and non-motor tasks are represented in posterior lobules (lobule VI – Crus I; Crus II – VIIB; lobules IX-X) [100].

In humans, limb and gait ataxia are more often associated with stroke involving the superior cerebellar artery (supplying blood flow to anterior lobules) as compared to the posterior inferior cerebellar artery (supplying blood flow to posterior lobules) [22–24]. Additionally, anterior lobules, including anterior parts of lobule VI, along with lobule VIII, receive spinal afferents through the spinocerebellar tracts [101]. These lobules are also reciprocally interconnected with the motor cortices [102]. Interestingly, individuals with infarction involving lobule VI, along with lobules VII-X, but sparing the anterior lobules, had a minor degree of motor impairment [25], further suggesting lobule VI’s dual role in non-motor and, to a lesser extent, motor behaviors. With respect to lesions of lobule IV/V in mice, impaired motor adaptation, gait coordination, and locomotor activity has been reported [26, 47]. In 16p11.2^dp/+^ mice, we found no gait or motor adaptation impairments as measured by the Erasmus Ladder, nor did we find any alterations in ectopic PC number or parvalbumin+ MLI cell counts relative to WT mice in lobule IV/V.

Conversely, stroke involving the posterior inferior cerebellar artery or damage to lobules VI, VII, and IX, results in cerebellar cognitive affective syndrome (CCAS), characterized by impairments in executive function, linguistic processing, spatial cognition, and affect regulation [27]. The behavioral phenotypes of CCAS originally led to the formation of Schmahmann’s dysmetria of thought hypothesis, which suggests that the cerebellum maintains both motor *and* cognitive behavior at a homeostatic baseline [103]. This hypothesis has also been applied to many of the neuropsychiatric disorders associated with the 16p11.2^dp/+^ CNV given their similar phenotypes. For example, it has been suggested that alterations in the cortico-thalamic-cerebellar circuit occur in SCZ and are associated with cognitive dysmetria [25, 104, 105]. Importantly, we have observed both behavioral and structural alterations in or associated with lobule VI in 16p11.2^dp/+^ mice, a posterior cerebellar lobule that has been associated with neuropsychiatric phenotypes and cognitive alterations in humans following damage. Further investigation of cognitive behaviors involving the cerebellum in 16p11.2^dp/+^ mice is crucial to identify possible links between the specific cerebellar structural abnormalities we observed.

### Behavioral phenotypes in humans with neuropsychiatric disorders associated with 16p11.2 microduplication

In most cases, our data mirror the behavioral phenotypes seen in the neuropsychiatric disorders associated with this CNV in humans. With respect to eyeblink conditioning, human patients with SCZ display reduced CR % [37–42]. Furthermore, longer CR onset latencies have been reported in SCZ patients [40, 43]. In eyeblink conditioning studies of children with ASD, typical trace eyeblink conditioning was reported, while altered delay eyeblink conditioning revealed shorter latency CRs [44], in contrast to our results from delay eyeblink conditioning in 16p11.2^dp/+^ mice. Additionally, indices of CRs tended to be lower in children with ADHD than in controls [106].

With respect to PPI, reduction in sensorimotor gating has been frequently reported in individuals with SCZ [60, 61]. PPI% reduction has also been reported in individuals with ASD compared to control groups [62]. Similarly, we observed that 16p11.2^dp/+^ mice exhibited deficits in PPI%, but this deficit was only observed in female 16p11.2^dp/+^ mice. This finding contrasts with some reports in the field, describing reduced PPI% in males with SCZ and not females [107]. However, another group using the same 16p11.2^dp/+^ mouse model in this study, found reduced PPI% in only female 16p11.2^dp/+^ mice as well [19], while others have reported typical sensorimotor gating in this 16p11.2^dp/+^ mouse model [59]. Taken together, the behavioral deficits in 16p11.2^dp/+^ mice resemble the behavioral deficits seen in humans with this CNV. Therefore, 16p11.2^dp/+^ mice display face validity to study the human etiology of neuropsychiatric disorders associated with this CNV and may be useful for studies that aim to test interventions for disorders associated with 16p11.2 microduplication.

## Conclusion

Our findings are consistent with previous studies implicating the cerebellum in motor learning, namely in that the 16p11.2^dp/+^ CNV did not structurally or behaviorally alter the motor-associated lobule IV/V, but it did alter lobule VI, a region recently linked to cognitive function [27]. However, further work is necessary to quantify other anterior and posterior lobule-associated behaviors, in addition to the examination of structural alterations in these lobules, to truly understand the extent of the cerebellum’s contributions to both motor and cognitive phenotypes in neuropsychiatric disorders associated with this CNV.

## ACKNOWLEDGEMENTS

The authors would like to thank Dr. Shane Heiney and the Neural Circuits and Behavior Core at the University of Iowa for training, behavioral equipment, and the confocal microscope used in this study. This project was funded by the Nellie Ball Collaborative and the Iowa Neuroscience Institute Carver Trust Schizophrenia Research Program of Excellence.

## REFERENCES

[1] Gillentine MA, Lupo PJ, Stankiewicz P, Schaaf CP. An estimation of the prevalence of genomic disorders using chromosomal microarray data. Journal of human genetics. 2018;63(7):795–801. doi:10.1038/s10038-018-0451-x

[2] Weiss LA, Shen Y, Korn JM, Arking DE, Miller DT, Fossdal R, Saemundsen E, Stefansson H, Ferreira MAR, Green T, et al. Association between microdeletion and microduplication at 16p11.2 and autism. The New England Journal of Medicine. 2008;358(7):667–675. doi:10.1056/NEJMoa075974

[3] McCarthy S, Makarov V, Kirov G, Addington A, McClellan J, Yoon S, Perkins D, Dickel D, Kusenda M, Krastoshevsky O, et al. Microduplications of 16p11.2 are associated with schizophrenia. Nat Genet. 2009;41(11):1223– 1227. doi:10.1038/ng.474

[4] Fernandez B, Roberts W, Chung B, Weksberg R, Meyn S, Szatmari P, Joseph-George A, Mackay S, Whitten K, Noble B, et al. Phenotypic spectrum associated with de novo and inherited deletions and duplications at 16p11.2 in individuals ascertained for diagnosis of autism spectrum disorder. J Med Genet. 2010;47(3):195–203. doi:10.1136/jmg.2009.069369

[5] Shinawi M, Liu P, Kang S-H, Shen J, Belmont J, Scott D, Probst F, Craigen W, Graham B, Pursley A, et al. Recurrent reciprocal 16p11.2 rearrangements associated with global developmental delay, behavioral problems, dysmorphism, epilepsy, and abnormal head size. J Med Genet. 2010;47(5):332–341. doi:10.1136/jmg.2009.073015

[6] Barber J, Hall V, Maloney V, Huang S, Roberts A, Brady A, Foulds N, Bewes B, Volleth M, Liehr T, et al. 16p11.2-p12.2 duplication syndrome; a genomic condition differentiated from euchromatic variation of 16p11.2. Eur J Hum Genet. 2013;21(2):182–189. doi:10.1038/ejhg.2012.144

[7] D’Angelo D, Lebon S, Chen Q, Martin-Brevet S, Green Snyder L, Hippolyte L, Hanson E, Maillard A, Faucett A, Macé A, et al. Defining the Effect of the 16p11.2 Duplication on Cognition, Behavior, and Medical Comorbidities. JAMA Psychiatry. 2016;73(1):20–30. doi:10.1001/jamapsychiatry.2015.2123

[8] Green Snyder L, D’Angelo D, Chen Q, Bernier R, Goin-Kochel R, Stevens Wallace A, Gerdts J, Kanne S, Berry L, Blaskey L, et al. Autism Spectrum Disorder, Developmental and Psychiatric Features in 16p11.2 Duplication. Journal of Autism and Developmental Disorders. 2016;46:2734–2748. 10.1007/s10803-016-2807-4

[9] Bernier R, Hudac C, Chen Q, Zeng C, Wallace AS, Gerdts J, Earl R, Peterson J, Wolken A, Peters A, et al. Developmental trajectories for young children with 16p11.2 copy number variation. American Journal of Medical genetics. Part B, Neuropsychiatric Genetics: the Official Publication of the International Society of Psychiatric Genetics. 2017;174(4):367–380. doi:10.1002/ajmg.b.32525

[10] Chang H, Lingyi L, Li M, Xiao X. Rare and common variants at 16p11.2 are associated with schizophrenia. Schizophrenia Research. 2017;184:105–108. doi:10.1016/j.schres.2016.11.031

[11] Sahoo T, Theisen A, Rosenfeld J, Lamb A, Ravnan B, Schultz R, Torchia B, Neill N, Casci I, Beijjani B, et al. Copy number variants of schizophrenia susceptibility loci are associated with a spectrum of speech and developmental delays and behavior problems. Genet Med. 2011;13(10):868–880. doi:10.1097/GIM.0b013e3182217a06

[12] Steinberg S, de Jong S, Mattheisen M, Costas J, Demontis D, Jamain S, Pietiläinen O, Lin K, Papiol S, Huttenlocher J, et al. Common variant at 16p11.2 conferring risk of psychosis. Mol Psychiatry. 2014;19(1):108–114. doi:10.1038/mp.2012.157

[13] Kirov G, Pocklington A, Holmans P, Ivanov D, Ikeda M, Ruderfer D, Moran J, Chambert K, Toncheva D, Georgieva L, et al. De novo CNV analysis implicates specific abnormalities of postsynaptic signalling complexes in the pathogenesis of schizophrenia. Mol Psychiatry. 2012;17(2):142–153. doi:10.1038/mp.2011.154

[14] Marshall CR, Howrigan DP, Merico D, Thiruvahindrapuram B, Wu W, Greer DS, Antaki D, Shetty A, Holmans PA, Pinto D, et al. Contribution of copy number variants to schizophrenia from a genome-wide study of 41,321 subjects. Nature Genetics. 2017;49(1):27–35. doi:10.1038/ng.3725

[15] Lin GN, Corominas R, Lemmens I, Yang X, Tavernier J, Hill DE, Vidal M, Sebat J, Iakoucheva LM. Spatiotemporal 16p11.2 protein network implicates cortical late mid-fetal brain development and KCTD13-Cul3-RhoA pathway in psychiatric diseases. Neuron. 2015;85(4):742–754. doi:10.1016/j.neuron.2015.01.010

[16] Rein B, Yan Z. 16p11.2 Copy Number Variations and Neurodevelopmental Disorders. Trends in Neurosciences. 2020;43(11):886–901. doi:10.1016/j.tins.2020.09.001

[17] Blackmon K, Thesen T, Green S, Ben-Avi E, Wang X, Fuchs B, Kuzniecky R, Devinsky O. Focal Cortical Anomalies and Language Impairment in 16p11.2 Deletion and Duplication Syndrome. Cerebral Cortex. 2018;28(7):2422–2430. doi:10.1093/cercor/bhx143

[18] Bertero A, Liska A, Pagani M, Parolisi R, Masferrer ME, Gritti M, Pedrazzoli M, Galbusera A, Sarica A, Cerasa A, et al. Autism-associated 16p11.2 microdeletion impairs prefrontal functional connectivity in mouse and human. Brain. 2018;141(7):2055–2065. doi:10.1093/brain/awy111

[19] Bristow GC, Thomson DM, Openshaw RL, Mitchell EJ, Pratt JA, Dawson N, Morris BJ. 16p11 Duplication Disrupts Hippocampal-Orbitofrontal-Amygdala Connectivity, Revealing a Neural Circuit Endophenotype for Schizophrenia. Cell Reports. 2020;31(3):107536. doi:10.1016/j.celrep.2020.107536

[20] Mega MS, Cummings JL. Frontal-subcortical circuits and neuropsychiatric disorders. The Journal of Neuropsychiatry and Clinical Neurosciences. 1994;6(4):358–370. doi:10.1176/jnp.6.4.358

[21] Tekin S, Cummings JL. Frontal-subcortical neuronal circuits and clinical neuropsychiatry: an update. Journal of Psychosomatic Research. 2002;53(2):647–654. doi:10.1016/s0022-3999(02)00428-2

[22] Kase CS, Norrving B, Levine SR, Babikian VL, Chodosh EH, Wolf PA, Welch KM. Cerebellar infarction. Clinical and anatomic observations in 66 cases. Stroke. 1993;24(1):76–83. doi:10.1161/01.STR.24.1.76

[23] Timmann D, Konczak J, Ilg W, Donchin O, Hermsdörfer J, Gizewski ER, Schoch B. Current advances in lesion-symptom mapping of the human cerebellum. Neuroscience. 2009;162(3):836–851. (New Insights in Cerebellar Function). doi:10.1016/j.neuroscience.2009.01.040

[24] Tohgi H, Takahashi S, Chiba K, Hirata Y. Cerebellar infarction. Clinical and neuroimaging analysis in 293 patients. The Tohoku Cerebellar Infarction Study Group. Stroke. 1993;24(11):1697–1701. doi:10.1161/01.STR.24.11.1697

[25] Jeremy D. Schmahmann. The cerebellar cognitive affective syndrome: clinical correlations of the dysmetria of thought hypothesis. International Review of Psychiatry. 2009. https://www.tandfonline.com/doi/abs/10.1080/09540260120082164

[26] Chao OY, Zhang H, Pathak SS, Huston JP, Yang Y-M. Functional Convergence of Motor and Social Processes in Lobule IV/V of the Mouse Cerebellum. Cerebellum (London, England). 2021 Mar 4. doi:10.1007/s12311-021-01246-7

[27] Jeremy D. Schmahmann. The cerebellum and cognition. Neuroscience Letters. 2018. doi:10.1016/j.neulet.2018.07.005

[28] Stoodley CJ, Schmahmann JD. Functional topography in the human cerebellum: A meta-analysis of neuroimaging studies. NeuroImage. 2009;44(2):489–501. doi:10.1016/j.neuroimage.2008.08.039

[29] Timmann D, Küper M, Gizewski ER, Schoch B, Donchin O. Lesion-Symptom Mapping of the Human Cerebellum. In: Manto MU, Gruol DL, Schmahmann JD, Koibuchi N, Sillitoe RV, editors. Handbook of the Cerebellum and Cerebellar Disorders. Cham: Springer International Publishing; 2022. p. 1857–1890. 10.1007/978-3-030-23810-0_72. doi:10.1007/978-3-030-23810-0_72

[30] Stoodley CJ, MacMore JP, Makris N, Sherman JC, Schmahmann JD. Location of lesion determines motor vs. cognitive consequences in patients with cerebellar stroke. NeuroImage: Clinical. 2016;12:765–775. doi:10.1016/j.nicl.2016.10.013

[31] Catherine J Stoodley, Jason P MacMore, Nikos Makris, Janet C Sherman, Jeremy D Schmahmann. Location of lesion determines motor vs. cognitive consequences in patients with cerebellar stroke. Neuroimage Clinical. 2016. doi:10.1016/j.nicl.2016.10.013

[32] Halverson HE, Kim J, Khilkevich A, Mauk MD, Augustine GJ. Feedback inhibition underlies new computational functions of cerebellar interneurons. eLife. 2022;11:e77603. doi:10.7554/eLife.77603

[33] Halverson HE, Khilkevich A, Mauk MD. Cerebellar Processing Common to Delay and Trace Eyelid Conditioning. The Journal of Neuroscience. 2018;38(33):7221–7236. doi:10.1523/JNEUROSCI.0430-18.2018

[34] Halverson HE, Khilkevich A, Mauk MD. Relating Cerebellar Purkinje Cell Activity to the Timing and Amplitude of Conditioned Eyelid Responses. Journal of Neuroscience. 2015;35(20):7813–7832. doi:10.1523/JNEUROSCI.3663-14.2015

[35] Perrett SP, Ruiz BP, Mauk MD. Cerebellar cortex lesions disrupt learning-dependent timing of conditioned eyelid responses. Journal of Neuroscience. 1993;13(4):1708–1718. doi:10.1523/JNEUROSCI.13-04-01708.1993

[36] Garcia KS, Steele PM, Mauk MD. Cerebellar Cortex Lesions Prevent Acquisition of Conditioned Eyelid Responses. Journal of Neuroscience. 1999;19(24):10940–10947. doi:10.1523/JNEUROSCI.19-24-10940.1999

[37] Hofer E, Doby D, Anderer P, Dantendorfer K. Impaired conditional discrimination learning in schizophrenia. Schizophrenia Research. 2001;51(2):127–136. doi:10.1016/S0920-9964(00)00118-3

[38] Brown SM, Kieffaber PD, Carroll CA, Vohs JL, Tracy JA, Shekhar A, O’Donnell BF, Steinmetz JE, Hetrick WP. Eyeblink conditioning deficits indicate timing and cerebellar abnormalities in schizophrenia. Brain and Cognition. 2005;58(1):94–108. (Neuropsychology of Timing and Time Perception). doi:10.1016/j.bandc.2004.09.011

[39] Edwards CR, Newman S, Bismark A, Skosnik PD, O’Donnell BF, Shekhar A, Steinmetz JE, Hetrick WP. Cerebellum volume and eyeblink conditioning in schizophrenia. Psychiatry Research: Neuroimaging. 2008;162(3):185–194. doi:10.1016/j.pscychresns.2007.06.001

[40] Bolbecker AR, Steinmetz AB, Mehta CS, Forsyth JK, Klaunig MJ, Lazar EK, Steinmetz JE, O’Donnell BF, Hetrick WP. Exploration of cerebellar-dependent associative learning in schizophrenia: Effects of varying and shifting interstimulus interval on eyeblink conditioning. Behavioral neuroscience. 2011;125(5):687–698. doi:10.1037/a0025150

[41] Forsyth JK, Bolbecker AR, Mehta CS, Klaunig MJ, Steinmetz JE, O’Donnell BF, Hetrick WP. Cerebellar-Dependent Eyeblink Conditioning Deficits in Schizophrenia Spectrum Disorders. Schizophrenia Bulletin. 2012;38(4):751–759. doi:10.1093/schbul/sbq148

[42] Coesmans M, Röder CH, Smit AE, Koekkoek SKE, Zeeuw CID, Frens MA, Geest JN van der. Cerebellar motor learning deficits in medicated and medication-free men with recent-onset schizophrenia. Journal of Psychiatry and Neuroscience. 2014;39(1):E3–E11. doi:10.1503/jpn.120205

[43] Marenco S, Weinberger DR, Schreurs BG. Single-cue delay and trace classical conditioning in schizophrenia. Biological Psychiatry. 2003;53(5):390–402. doi:10.1016/S0006-3223(02)01506-8

[44] Oristaglio J, West SH, Ghaffari M, Lech MS, Verma BR, Harvey JA, Welsh JP, Malone RP. Children with autism spectrum disorders show abnormal conditioned response timing on delay, but not trace, eyeblink conditioning. Neuroscience. 2013;248:708–718. doi:10.1016/j.neuroscience.2013.06.007

[45] Giessen RSVD, Koekkoek SK, Dorp S van, Gruijl JRD, Cupido A, Khosrovani S, Dortland B, Wellershaus K, Degen J, Deuchars J, et al. Role of Olivary Electrical Coupling in Cerebellar Motor Learning. Neuron. 2008;58(4):599–612. doi:10.1016/j.neuron.2008.03.016

[46] Schonewille M, Gao Z, Boele H-J, Veloz MFV, Amerika WE, Simek AAM, De Jeu MT, Steinberg JP, Takamiya K, Hoebeek FE, et al. Reevaluating the Role of LTD in Cerebellar Motor Learning. Neuron. 2011;70(1):43–50. doi:10.1016/j.neuron.2011.02.044

[47] Liu X-X, Chen X-H, Zheng Z-W, Jiang Q, Li C, Yang L, Chen X, Mao X-F, Yuan H-Y, Feng L-L, et al. BOD1 regulates the cerebellar IV/V lobe-fastigial nucleus circuit associated with motor coordination. Signal Transduction and Targeted Therapy. 2022;7(1):1–14. doi:10.1038/s41392-022-00989-x

[48] Peter S, ten Brinke MM, Stedehouder J, Reinelt CM, Wu B, Zhou H, Zhou K, Boele H-J, Kushner SA, Lee MG, et al. Dysfunctional cerebellar Purkinje cells contribute to autism-like behaviour in Shank2-deficient mice. Nature Communications. 2016;7:12627. doi:10.1038/ncomms12627

[49] Palmen SJMC, van Engeland H, Hof PR, Schmitz C. Neuropathological findings in autism. Brain. 2004;127(12):2572– 2583. doi:10.1093/brain/awh287

[50] de la Torre-Ubieta L, Won H, Stein JL, Geschwind DH. Advancing the understanding of autism disease mechanisms through genetics. Nature Medicine. 2016;22(4):345–361. doi:10.1038/nm.4071

[51] Wegiel J, Flory M, Kuchna I, Nowicki K, Ma SY, Imaki H, Wegiel J, Cohen IL, London E, Wisniewski T, et al. Stereological study of the neuronal number and volume of 38 brain subdivisions of subjects diagnosed with autism reveals significant alterations restricted to the striatum, amygdala and cerebellum. Acta Neuropathologica Communications. 2014;2(1):141. doi:10.1186/s40478-014-0141-7

[52] Fehlow P, Bernstein K, Tennstedt A, Walther F. [Early infantile autism and excessive aerophagy with symptomatic megacolon and ileus in a case of Ehlers-Danlos syndrome]. Padiatrie und Grenzgebiete. 1993;31(4):259–267.

[53] Maloku E, Covelo IR, Hanbauer I, Guidotti A, Kadriu B, Hu Q, Davis JM, Costa E. Lower number of cerebellar Purkinje neurons in psychosis is associated with reduced reelin expression. Proceedings of the National Academy of Sciences of the United States of America. 2010;107(9):4407–4411. doi:10.1073/pnas.0914483107

[54] Mavroudis IA, Petrides F, Manani M, Chatzinikolaou F, Ciobică AS, Pădurariu M, Kazis D, Njau SN, Costa VG, Baloyannis SJ. Purkinje cells pathology in schizophrenia. A morphometric approach. Romanian Journal of Morphology and Embryology = Revue Roumaine De Morphologie Et Embryologie. 2017;58(2):419–424.

[55] Qureshi AY, Mueller S, Snyder AZ, Mukherjee P, Berman JI, Roberts TPL, Nagarajan SS, Spiro JE, Chung WK, Sherr EH, et al. Opposing Brain Differences in 16p11.2 Deletion and Duplication Carriers. The Journal of Neuroscience. 2014;34(34):11199–11211. doi:10.1523/JNEUROSCI.1366-14.2014

[56] GTEx Consortium. The GTEx Consortium atlas of genetic regulatory effects across human tissues. Science (New York, N.Y.). 2020;369(6509):1318–1330. doi:10.1126/science.aaz1776

[57] Horev G, Ellegood J, Lerch J, Son Y-E, Muthuswamy L, Vogel H, Krieger A, Buja A, Henkelman M, Wigler M, et al. Dosage-Dependent Phenotypes in Models of Human 16p11.2 Lesions Found in Autism. Proceedings of the National Academy of Sciences of the United States of America. 2011;108(41):17076–17081. doi:10.1073/pnas.1114042108

[58] Tai DJC, Razaz P, Erdin S, Gao D, Wang J, Nuttle X, de Esch CE, Collins RL, Currall BB, O’Keefe K, et al. Tissue– and cell-type-specific molecular and functional signatures of 16p11.2 reciprocal genomic disorder across mouse brain and human neuronal models. American Journal of Human Genetics. 2022;109(10):1789–1813. doi:10.1016/j.ajhg.2022.08.012

[59] Rein B, Tan T, Yang F, Wang W, Williams J, Zhang F, Mills A, Yan Z. Reversal of synaptic and behavioral deficits in a 16p11.2 duplication mouse model via restoration of the GABA synapse regulator Npas4. Molecular Psychiatry. 2021;26(6):1967–1979. doi:10.1038/s41380-020-0693-9

[60] Mena A, Ruiz-Salas JC, Puentes A, Dorado I, Ruiz-Veguilla M, De la Casa LG. Reduced Prepulse Inhibition as a Biomarker of Schizophrenia. Frontiers in Behavioral Neuroscience. 2016;10(202). doi:10.3389/fnbeh.2016.00202

[61] San-Martin R, Castro LA, Menezes PR, Fraga FJ, Simões PW, Salum C. Meta-Analysis of Sensorimotor Gating Deficits in Patients With Schizophrenia Evaluated by Prepulse Inhibition Test. Schizophrenia Bulletin. 2020;46(6):1482–1497. doi:10.1093/schbul/sbaa059

[62] Cheng C-H, Chan P-YS, Hsu S-C, Liu C-Y. Meta-analysis of sensorimotor gating in patients with autism spectrum disorders. Psychiatry Research. 2018;262:413–419. doi:10.1016/j.psychres.2017.09.016

[63] Lauffer M, Wen H, Myers B, Plumb A, Parker K, Williams A. Deletion of the voltage-gated calcium channel, CaV1.3, causes deficits in motor performance and associative learning. Genes, Brain and Behavior. 2022;21(2):e12791. doi:10.1111/gbb.12791

[64] Brown AM, Arancillo M, Lin T, Catt DR, Zhou J, Lackey EP, Stay TL, Zuo Z, White JJ, Sillitoe RV. Molecular layer interneurons shape the spike activity of cerebellar Purkinje cells. Scientific Reports. 2019;9(1):1742. doi:10.1038/s41598-018-38264-1

[65] Heiney SA, Ohmae S, Kim OA, Medina JF. Single-Unit Extracellular Recording from the Cerebellum During Eyeblink Conditioning in Head-Fixed Mice. In: Sillitoe RV, editor. Extracellular Recording Approaches. New York, NY: Springer; 2018. p. 39–71. (Neuromethods). 10.1007/978-1-4939-7549-5_3. doi:10.1007/978-1-4939-7549-5_3

[66] Jacquemont S, Reymond A, Zufferey F, Harewood L, Walters RG, Kutalik Z, Martinet D, Shen Y, Valsesia A, Beckmann ND, et al. Mirror extreme BMI phenotypes associated with gene dosage at the chromosome 16p11.2 locus. Nature. 2011;478(7367):97–102. doi:10.1038/nature10406

[67] ten Brinke MM, Boele H-J, Spanke JK, Potters J-W, Kornysheva K, Wulff P, IJpelaar ACHG, Koekkoek SKE, De Zeeuw CI. Evolving Models of Pavlovian Conditioning: Cerebellar Cortical Dynamics in Awake Behaving Mice. Cell Reports. 2015;13(9):1977–1988. doi:10.1016/j.celrep.2015.10.057

[68] Braff DL, Geyer MA, Swerdlow NR. Human studies of prepulse inhibition of startle: normal subjects, patient groups, and pharmacological studies. Psychopharmacology. 2001;156(2–3):234–258. doi:10.1007/s002130100810

[69] Diwakar S, Lombardo P, Solinas S, Naldi G, D’Angelo E. Local Field Potential Modeling Predicts Dense Activation in Cerebellar Granule Cells Clusters under LTP and LTD Control. PLoS ONE. 2011;6(7):e21928. doi:10.1371/journal.pone.0021928

[70] Sillitoe RV, Gopal N, Joyner AL. Embryonic origins of ZebrinII parasagittal stripes and establishment of topographic Purkinje cell projections. Neuroscience. 2009;162(3):574–588. (New Insights in Cerebellar Function). doi:10.1016/j.neuroscience.2008.12.025

[71] Palmer LM, Clark BA, Gründemann J, Roth A, Stuart GJ, Häusser M. Initiation of simple and complex spikes in cerebellar Purkinje cells. The Journal of Physiology. 2010;588(Pt 10):1709–1717. doi:10.1113/jphysiol.2010.188300

[72] Barmack NH, Yakhnitsa V. Functions of Interneurons in Mouse Cerebellum. The Journal of Neuroscience. 2008;28(5):1140–1152. doi:10.1523/jneurosci.3942-07.2008

[73] Badura A, Verpeut JL, Metzger JW, Pereira TD, Pisano TJ, Deverett B, Bakshinskaya DE, Wang SS-H. Normal cognitive and social development require posterior cerebellar activity. eLife. 2018;7:e36401. doi:10.7554/eLife.36401

[74] Chae T, Kwon YT, Bronson R, Dikkes P, Li E, Tsai L-H. Mice Lacking p35, a Neuronal Specific Activator of Cdk5, Display Cortical Lamination Defects, Seizures, and Adult Lethality. Neuron. 1997;18(1):29–42. doi:10.1016/S0896-6273(01)80044-1

[75] Goffinet AM. The embryonic development of the cerebellum in normal and reeler mutant mice. Anatomy and Embryology. 1983;168(1):73–86. doi:10.1007/BF00305400

[76] Yuasa S, Kitoh J, Oda S, Kawamura K. Obstructed migration of Purkinje cells in the developing cerebellum of the reeler mutant mouse. Anatomy and Embryology. 1993;188(4):317–329. doi:10.1007/BF00185941

[77] Miyata T, Ono Y, Okamoto M, Masaoka M, Sakakibara A, Kawaguchi A, Hashimoto M, Ogawa M. Migration, early axonogenesis, and Reelin-dependent layer-forming behavior of early/posterior-born Purkinje cells in the developing mouse lateral cerebellum. Neural Development. 2010;5(1):1–22. doi:10.1186/1749-8104-5-23

[78] Chung S, Zhang Y, Van Der Hoorn F, Hawkes R. The anatomy of the cerebellar nuclei in the normal and scrambler mouse as revealed by the expression of the microtubule-associated protein kinesin light chain 3. Brain Research. 2007;1140:120–131. (A Catalog of the Neurological Mutants of the Mouse Revisited). doi:10.1016/j.brainres.2006.01.100

[79] Chung S-H, Sillitoe RV, Croci L, Badaloni A, Consalez G, Hawkes R. Purkinje cell phenotype restricts the distribution of unipolar brush cells. Neuroscience. 2009;164(4):1496–1508. doi:10.1016/j.neuroscience.2009.09.080

[80] Reddy SS, Connor TE, Weeber EJ, Rebeck W. Similarities and differences in structure, expression, and functions of VLDLR and ApoER2. Molecular Neurodegeneration. 2011;6(1):30. doi:10.1186/1750-1326-6-30

[81] Kuo G, Arnaud L, Kronstad-O’Brien P, Cooper JA. Absence of Fyn and Src Causes a Reeler-Like Phenotype. Journal of Neuroscience. 2005;25(37):8578–8586. doi:10.1523/JNEUROSCI.1656-05.2005

[82] Hadj-Sahraoui N, Frederic F, Zanjani H, Herrup K, Delhaye-Bouchaud N, Mariani J. Purkinje cell loss in heterozygous staggerer mutant mice during aging. Developmental Brain Research. 1997;98(1):1–8. doi:10.1016/S0165-3806(96)00153-8

[83] Yue Q, Groszer M, Gil JS, Berk AJ, Messing A, Wu H, Liu X. PTEN deletion in Bergmann glia leads to premature differentiation and affects laminar organization. Development. 2005;132(14):3281–3291. doi:10.1242/dev.01891

[84] Armengol J-A, Sotelo C. Early dendritic development of Purkinje cells in the rat cerebellum. A light and electron microscopic study using axonal tracing in ‘in vitro’ slices. Developmental Brain Research. 1991;64(1):95–114. doi:10.1016/0165-3806(91)90213-3

[85] Dey J, Ditzler S, Knoblaugh SE, Hatton BA, Schelter JM, Cleary MA, Mecham B, Rorke-Adams LB, Olson JM. A Distinct Smoothened Mutation Causes Severe Cerebellar Developmental Defects and Medulloblastoma in a Novel Transgenic Mouse Model. Molecular and Cellular Biology. 2012;32(20):4104–4115. doi:10.1128/MCB.00862-12

[86] Ma Q, Jones D, Borghesani PR, Segal RA, Nagasawa T, Kishimoto T, Bronson RT, Springer TA. Impaired B-lymphopoiesis, myelopoiesis, and derailed cerebellar neuron migration in CXCR4– and SDF-1-deficient mice. Proceedings of the National Academy of Sciences. 1998;95(16):9448–9453. doi:10.1073/pnas.95.16.9448

[87] Larouche M, Hawkes R. From clusters to stripes: The developmental origins of adult cerebellar compartmentation. The Cerebellum. 2006;5(2):77–88. doi:10.1080/14734220600804668

[88] Huang G-J, Edwards A, Tsai C-Y, Lee Y-S, Peng L, Era T, Hirabayashi Y, Tsai C-Y, Nishikawa S-I, Iwakura Y, et al. Ectopic Cerebellar Cell Migration Causes Maldevelopment of Purkinje Cells and Abnormal Motor Behaviour in Cxcr4 Null Mice. PLOS ONE. 2014;9(2):e86471. doi:10.1371/journal.pone.0086471

[89] M. Lafarga; M.T. Berciano; M. Blanco. Ectopic Purkinje Cells in the Cerebellar White Matter of Normal Adult Rodents: A Golgi Study. Acta Anatomica. 1987;127(1):53–58. 10.1159/000146237

[90] Vermeren MM, Zhang Q, Smethurst E, Segonds-Pichon A, Schrewe H, Wakelam MJO. The Phospholipase D2 Knock Out Mouse Has Ectopic Purkinje Cells and Suffers from Early Adult-Onset Anosmia. PLOS ONE. 2016;11(9):e0162814. doi:10.1371/journal.pone.0162814

[91] Butts T, Green MJ, Wingate RJT. Development of the cerebellum: simple steps to make a ‘little brain.’ Development. 2014;141(21):4031–4041. doi:10.1242/dev.106559

[92] Willis A, Pratt JA, Morris BJ. BDNF and JNK Signaling Modulate Cortical Interneuron and Perineuronal Net Development: Implications for Schizophrenia-Linked 16p11.2 Duplication Syndrome. Schizophrenia Bulletin. 2021;47(3):812–826. doi:10.1093/schbul/sbaa139

[93] Deshpande A, Yadav S, Dao DQ, Wu Z-Y, Hokanson KC, Cahill MK, Wiita AP, Jan Y-N, Ullian EM, Weiss LA. Cellular Phenotypes in Human iPSC-Derived Neurons from a Genetic Model of Autism Spectrum Disorder. Cell Reports. 2017;21(10):2678–2687. doi:10.1016/j.celrep.2017.11.037

[94] Filice F, Vörckel KJ, Sungur AÖ, Wöhr M, Schwaller B. Reduction in parvalbumin expression not loss of the parvalbumin-expressing GABA interneuron subpopulation in genetic parvalbumin and shank mouse models of autism. Molecular Brain. 2016;9(1):10. doi:10.1186/s13041-016-0192-8

[95] Lodge DJ, Behrens MM, Grace AA. A loss of parvalbumin-containing interneurons is associated with diminished oscillatory activity in an animal model of schizophrenia. The Journal of Neuroscience: The Official Journal of the Society for Neuroscience. 2009;29(8):2344–2354. doi:10.1523/JNEUROSCI.5419-08.2009

[96] Hashimoto T, Volk DW, Eggan SM, Mirnics K, Pierri JN, Sun Z, Sampson AR, Lewis DA. Gene Expression Deficits in a Subclass of GABA Neurons in the Prefrontal Cortex of Subjects with Schizophrenia. Journal of Neuroscience. 2003;23(15):6315–6326. doi:10.1523/JNEUROSCI.23-15-06315.2003

[97] Cooper JA. A mechanism for inside-out lamination in the neocortex. Trends in Neurosciences. 2008;31(3):113–119. doi:10.1016/j.tins.2007.12.003

[98] Xavier Guell & Jeremy Schmahmann. Cerebellar Functional Anatomy: a Didactic Summary Based on Human fMRI Evidence. The cerebellum. 2020. 10.1007/s12311-019-01083-9

[99] O. Oscarsson. Functional Organization of the Spino– and Cuneocerebellar Tracts. 1965. https://journals.physiology.org/doi/abs/10.1152/physrev.1965.45.3.495

[100] Brodal P. THE CORTICOPONTINE PROJECTION IN THE RHESUS MONKEY ORIGIN AND PRINCIPLES OF ORGANIZATION. Brain. 1978;101(2):251–283. doi:10.1093/brain/101.2.251

[101] Schmahmann JD. The cerebellum and cognition. Neuroscience Letters. 2019;688: 62–75. doi:10.1016/j.neulet.2018.07.005

[102] Andreasen NC, Paradiso S, O’Leary DS. “Cognitive dysmetria” as an integrative theory of schizophrenia: a dysfunction in cortical-subcortical-cerebellar circuitry? Schizophrenia Bulletin. 1998;24(2):203–218. doi:10.1093/oxfordjournals.schbul.a033321

[103] Andreasen NC, Nopoulos P, O’Leary DS, Miller DD, Wassink T, Flaum M. Defining the phenotype of schizophrenia: cognitive dysmetria and its neural mechanisms. Biological Psychiatry. 1999;46(7):908–920. doi:10.1016/S0006-3223(99)00152-3

[104] Frings M, Gaertner K, Buderath P, Gerwig M, Christiansen H, Schoch B, Gizewski ER, Hebebrand J, Timmann D. Timing of conditioned eyeblink responses is impaired in children with attention-deficit/hyperactivity disorder. Experimental Brain Research. 2010;201(2):167–176. doi:10.1007/s00221-009-2020-1

[105] Kumari V, Aasen I, Sharma T. Sex differences in prepulse inhibition deficits in chronic schizophrenia. Schizophrenia Research. 2004;69(2–3):219–235. doi:10.1016/j.schres.2003.09.010

